# Correlated substitutions reveal SARS-like coronaviruses recombine frequently with a diverse set of structured gene pools

**DOI:** 10.1101/2022.08.26.505425

**Authors:** Asher Preska Steinberg, Olin K. Silander, Edo Kussell

## Abstract

Quantifying SARS-like coronavirus (SL-CoV) evolution is critical to understanding the origins of SARS-CoV-2 and the molecular processes that could underlie future epidemic viruses. While genomic evidence implicates recombination as a factor in the emergence of SARS-CoV-2, few studies have quantified recombination rates among SL-CoVs. Here, we infer recombination rates of SL-CoVs from correlated substitutions in sequencing data using a coalescent model with recombination. Our computationally-efficient, non-phylogenetic method infers recombination parameters of both sampled sequences and the unsampled gene pools with which they recombine. We apply this approach to infer recombination parameters for a range of positive-sense RNA viruses. We then analyze a set of 191 SL-CoV sequences (including SARS-CoV-2) and find that ORF1ab and S genes frequently undergo recombination. We identify which SL-CoV sequence clusters have recombined with shared gene pools, and show that these pools have distinct structures and high recombination rates, with multiple recombination events occurring per synonymous substitution. We find that individual genes have recombined with different viral reservoirs. By decoupling contributions from mutation and recombination, we recover the phylogeny of non-recombined portions for many of these SL-CoVs, including the position of SARS-CoV-2 in this clonal phylogeny. Lastly, by analyzing 444,145 SARS-CoV-2 whole genome sequences, we show current diversity levels are insufficient to infer the within-population recombination rate of the virus since the pandemic began. Our work offers new methods for inferring recombination rates in RNA viruses with implications for understanding recombination in SARS-CoV-2 evolution and the structure of clonal relationships and gene pools shaping its origins.

**Significance Statement:** Quantifying the population genetics of SARS-like coronavirus (SL-CoV) evolution is vital to deciphering the origins of SARS-CoV-2 and pinpointing viruses with epidemic potential. While some Bayesian approaches can quantify recombination for these pathogens, the required simulations of recombination networks do not scale well with the massive amounts of sequences available in the genomics era. Our approach circumvents this by measuring correlated substitutions in sequences and fitting these data to a coalescent model with recombination. This allows us to analyze hundreds of thousands of sample sequences, and infer recombination rates for unsampled viral reservoirs. Our results provide insights into both the clonal relationships of sampled SL-CoV sequence clusters and the evolutionary dynamics of the gene pools with which they recombine.

## Introduction

Recombination can enable viruses to rapidly adapt to selective pressures (1–4) and to avoid accumulation of deleterious mutations that can lead to viral decline and extinction (5–7). Positive-sense single stranded RNA ((+)ssRNA) viruses display highly variable levels of recombination (8, 9), with some species such as *West Nile* and *Yellow fever viruses* showing scant evidence of recombination (10) and others such as those of the *Coronaviridae* family showing evidence of frequent recombination (11). During the ongoing COVID-19 pandemic, population genomics has played an invaluable role in tracking the spread of SARS-CoV-2 and its variants (12–14), as well as understanding correlations between genomic substitutions and transmission patterns (15–19). However, a quantitative, population genomics-based understanding of the relative contributions of recombination and mutation to the evolution of SARS-CoV-2 and other SARS-like coronaviruses is still being developed (20–27). Such knowledge will be important to understanding the emergence of past and future viruses at the source of major epidemics.

The majority of tools for studying recombination in RNA viruses are phylogeny-based, where recombination breakpoints are assessed by examining phylogenetic incongruence and Bayesian and Markov chain Monte Carlo techniques are used to infer recombination parameters (20–25, 28). These approaches have been successful at identifying instances of recombination, yet their application to large-scale population genomics data remains challenging due to the computational demands of these methods. Importantly, the inferred recombination parameters rely only on the observed (i.e., sampled) sequences, while recombination within the much larger, unobserved gene pools with which these branches interact is not captured by these models. Here, to infer the recombination parameters of (+)ssRNA viruses, we adapt our non-phylogenetic, computationally-efficient *mcorr* method, which we originally developed to measure homologous recombination rates in bacteria (29–31). In contrast to previous approaches which focus on recombination within sampled sequences, we infer recombination parameters for both sampled sequences and the larger gene pools they recombine with, revealing that SARS-like coronaviruses recombine with a diverse set of gene pools which have high levels of recombination.

## Results

### Using correlated substitutions to infer recombination rates in RNA viruses

Our primary aim is to infer the mutation and recombination rate within a set of sampled viral sequences, as well as the diversity and recombination rate of the unsampled pool of viruses with which the sampled viruses recombine. We model a ‘sample’ of viral lineages with mean coalescence time 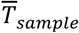, which replicate, mutate (at rate *μ*), and recombine (at rate *γ*) with a much larger ‘pool’ of lineages that have a mean coalescence time 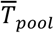 (Fig. 1A). We originally applied this model to infer recombination rates for bacteria (30), with the key difference here being introduced by the process of ‘copy-choice’ recombination in RNA viruses (described in more detail below). The model predicts the conditional probability of a synonymous substitution at a genomic site *i* + *l* given a substitution at site *i*, which we refer to as the ‘correlation profile’, *P*(*l*), where *l* is the distance between sites in nucleotides (nt). In a highly recombining viral population, this profile should decline rapidly as the distance between sites increases, while in a non-recombining population, this profile should be largely flat (see Fig. 1B for schematic) (29). To measure correlation profiles, we use sets of whole genome sequences (WGS) as our samples, and use alignments of coding regions (CDS) to determine synonymous substitutions for all possible sequence pairs. For a sequence pair within the sample, we assign a binary variable σ_*i*_ a value of 1 for a substitution and a 0 for identity at position *i* (we refer to *σ_i_* as the substitution profile). We exclusively consider third-position codon sites which are fourfold degenerate. The correlation profile is obtained by *P*(*l*) ≡ *P*(*σ_i+l_*|*σ_i_* = 1), where the conditional probability is computed over all possible sequences pairs and averaged over all positions *i*.

**Figure 1.**
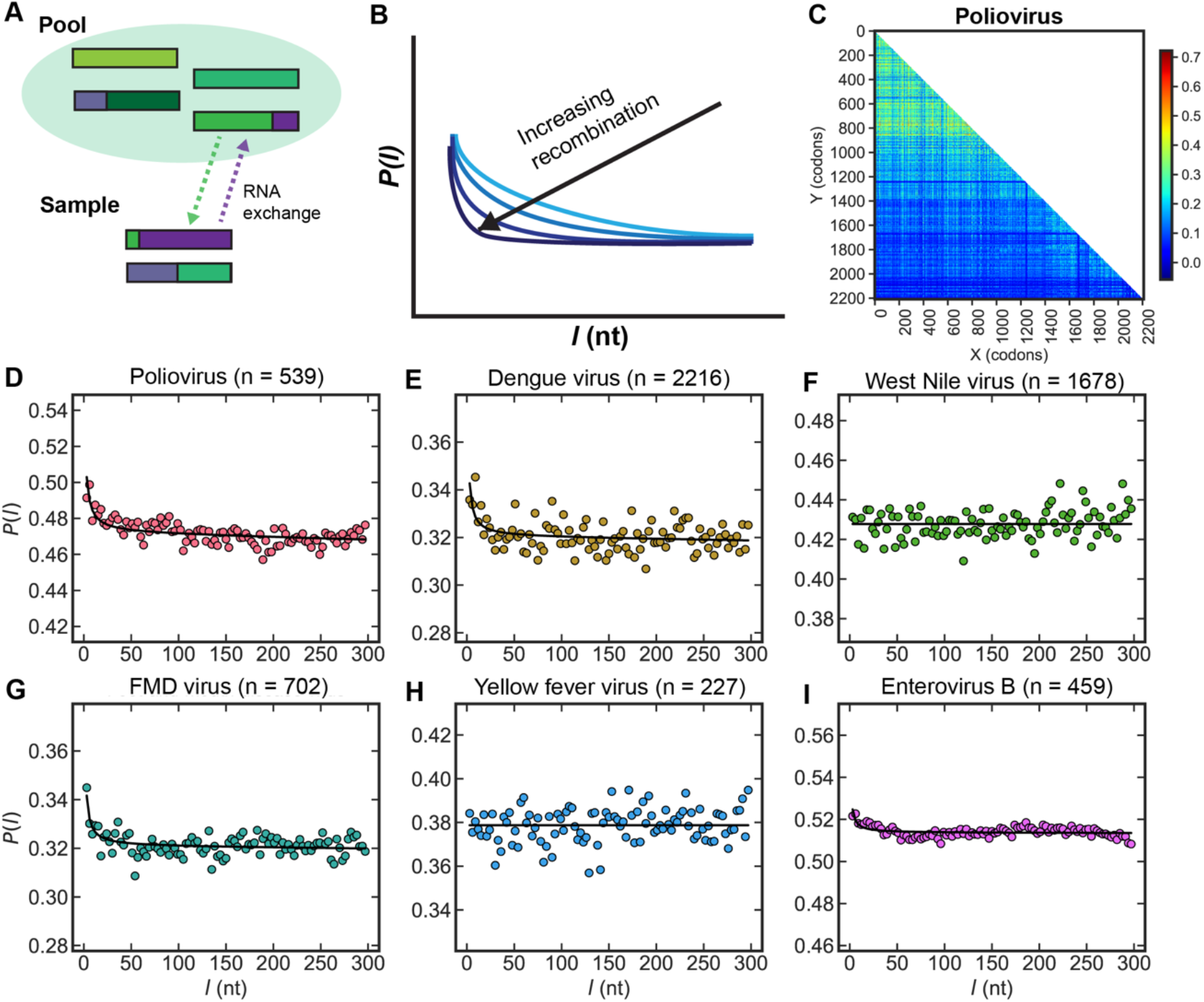
Correlated substitutions in RNA viruses. (A) Schematic depicting the exchange of homologous RNA between a set of lineages (i.e., the “sample”) and a larger “pool” of lineages via a copy-choice mechanism. (B) Schematic depicting different correlation profiles of synonymous substitutions with various levels of recombination. (C) A heatmap of Pearson’s correlation coefficient (*ρ*(*X, Y*)) of synonymous substitutions across the coding region of the *Poliovirus* genome calculated using 539 *Poliovirus* genomes (see Materials and Methods for more details). Each position (*X, Y*) displays the corresponding value of *ρ*(*X, Y*) as a color for a pair of codons located at genomic positions *X* and *Y* (given in codons). Color bar indicates the value of *ρ*(*X,Y*). Monomorphic sites are assigned *ρ*(*X,Y*) = 0. (D-I) Correlation profiles of synonymous substitutions for a range of (+)ssRNA viruses. Markers correspond to the correlation profile *P*(*l*) for a given separation distance *l* (given in nucleotides, *nt*). The fit is shown as a solid line, and was performed under the assumption that only complete RNA strands are exchanged during template-switching events (i.e., we used the “template-switching model” described in Materials and Methods and the main text). Model selection was performed with the Akaike Information Criterion (AIC) to determine if a coalescent model with or without recombination best fit the data (see Materials and Methods for details). Parameters of homologous recombination are given in Table 1. *n* is the number of whole genome sequences analyzed. “FMD” virus stands for “Foot-and-mouth-disease virus”.

The model has two free parameters which are determined by fitting the predicted *P*(*l*) to its measured values from viral genome sequences. From the fit, we then calculate several useful quantities, including the pool’s mutational and recombinational divergence (given by 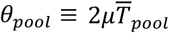 and 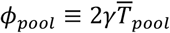) which respectively correspond to the expected number of synonymous substitution and recombination events per site since coalescence of the pool; the sample’s recombination coverage (*c_sample_*), which is the fraction of sites that recombined since coalescence of the sample; the sample’s mutational divergence 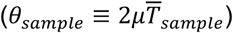, which is proportional to the average age of the clonal (i.e., non-recombined) genomic portions of the sample; and the relative recombination rate of the pool (*ϕ_pool_/θ_pool_* = *γ/μ*) which we denote as (*γ/μ*)_*pool*_. Unlike the synonymous substitutions, which we assume to be largely neutral, recombination events may be due to selective pressure or neutral drift.

Recombination in RNA viruses is thought to occur via a ‘copy-choice’ mechanism, in which RNA-dependent RNA polymerase (RdRP) switches from one RNA template to another during RNA synthesis while remaining bound to the nascent RNA strand, creating RNA with hybrid ancestry (8, 9, 32). Here, we focus on copy-choice recombination resulting in homologous recombination; non-homologous or ‘illegitimate’ recombination (i.e., insertion/deletion events) has previously been studied in SARS-like coronaviruses (23, 33) and RNA viruses (8) and is thought to be comparatively rare. This template-switching process occurs in the model at rate *γ* per site per viral replication (i.e., generation), and yields a hybrid viral genome consisting of a left arm from one genome and a right arm from another genome, joined at the recombination breakpoint (see Fig. S1a). Experiments performed with murine hepatitis virus (a *betacoronavirus*) indicate that recombination can occur during negative or positive strand synthesis, and that template-switching events do not exclusively occur when two live viruses co-infect a cell but can also occur with transfected RNA fragments as small as 450 nucleotides when RdRP switches from template to fragment (or vice versa) during synthesis (34). This latter scenario is similar to homologous recombination in bacteria, where DNA fragments of average size 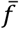 are taken up by the cell at rate *γ* per site per generation and incorporated within a genome, replacing the homologous sites (see Fig. S1b). In both the fragment incorporation model and the template-switching model, the predicted form of *P*(*l*) depends on the total recombination rate at a given locus (*r*), with 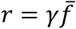 in the fragment-incorporation model and *r* = *γL* in the template-switching model, where *L* is the size of the genome (see Methods for functional forms). We note that the fragment-incorporation model has an extra fitting parameter 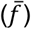.

We first analyzed correlated substitutions in *Poliovirus*, as this virus is known to have undergone substantial recombination during its evolution (9, 35–38). A genome-wide plot of the Pearson’s correlation coefficient for all pairwise synonymous substitutions (which is the square root of the classic linkage disequilibrium metric ‘*r*^2^’ but uses paired differences instead of allelic values (39)) across the CDS region of all major serotypes of *Poliovirus* (539 whole genome sequences used) shows that while substitutions tend to be more strongly correlated in the first ~800 codons, statistically significant correlations are found across the entire genome (Fig. 1C; see Material and Methods for more details of calculation). Fitting the correlation profile for *Poliovirus*, we inferred the parameters of homologous recombination (Fig. 1D, Table 1). To estimate the range of these parameters, we calculated 95% bootstrap confidence intervals by sampling the 539 genomes with replacement to create bootstrap replicates (Table 1; see Materials and Methods for details). Consistent with the literature, we found that *Poliovirus* has recombined substantially. We then proceeded to compute correlation profiles and infer recombination parameters for 12 other (+)ssRNA viruses (Fig. 1E-I, Fig. S2; Table 1, S1). We found results consistent with the literature, with viruses known to recombine showing evidence of recombination, e.g., *Dengue virus, Foot-and-mouth-disease virus*, and *Enterovirus B* (9, 10, 40–49), while others where little or no recombination has been reported such as *West Nile* and *Yellow fever virus* (10, 50, 51) did not show signatures of recombination. We fit the correlation profiles in Fig. 1 using the template-switching model (Table 1) or the fragment-incorporation model (Table S2), and found similar results; in particular, the predicted mean fragment size is generally on the order of the genome size and model selection does not strongly favor one model over the other. We therefore use the two-parameter template-switching model in all our analyses below.

**Table 1.** Parameters of homologous recombination for viruses shown in main text. If the entire genome was fit, the gene is listed as “full genome”. “SL-CoV” refers to the SARS-like betacoronaviruses. “orf1a” refers to the orf1ab CDS region before the −1 ribosomal frameshift and “orf1b” refers to the region after this frameshift. Parameters are given as the values inferred from the data, followed by the 95% bootstrap CI in square brackets (see Materials and Methods for calculation). We used model selection with AIC to determine if the profile was better fit with a coalescent model with or without recombination (see Materials and Methods). For those profiles which were better fit with the coalescent model with recombination, we assumed that only template-switching occurs (i.e., we used the “template-switching model” described in Materials and Methods). For those profiles better fit by the model without recombination, coverage was set to c = 0 and no bootstrapping was performed. *L* is the length of the genome in the template-switching model, *w_t_/w_z_* is the Akaike weight of the template switching model (*w_t_*) over the weight of the model without recombination (*w_z_*; see Materials and Methods). All other parameters are described in main text.

| virus | gene | # of<br>seqs | $d_{sample}$ | $\theta_{pool}$ | $\phi_{pool}$ | $(\gamma/\mu)_{pool}$ | $L$ (nt) | evidence<br>ratio<br>($w_t/w_z$) |
| --- | --- | --- | --- | --- | --- | --- | --- | --- |
| Poliovirus | full<br>genome | 539 | 0.369<br>[0.359, 0.376] | 1.29<br>[1.25, 1.29] | 2.68<br>[2.42, 2.99] | 2.11<br>[1.93, 2.32] | 7.50E+03 | 9.96E+14 |
| Dengue virus | full<br>genome | 2216 | 0.256<br>[0.252, 0.260] | 0.559<br>[0.543, 0.575] | 2.93<br>[2.82, 3.04] | 5.24<br>[4.97, 5.57] | 1.10E+04 | 2.78E+04 |
| West Nile virus | full<br>genome | 1678 | 0.109 | n/a | n/a | n/a | n/a | 1.14E-06 |
| Foot-and-mouth<br>disease virus | full<br>genome | 702 | 0.290<br>[0.284, 0.295] | 0.560<br>[0.548, 0.572] | 3.21<br>[3.02, 3.41] | 5.73<br>[5.37, 6.14] | 8.30E+03 | 1.68E+08 |
| Yellow fever<br>virus | full<br>genome | 227 | 0.224 | n/a | n/a | n/a | n/a | 7.54E-03 |
| Enterovirus B | full<br>genome | 459 | 0.509<br>[0.506, 0.509] | 1.63<br>[1.61, 1.64] | 10.2<br>[9.37, 11.1] | 6.25<br>[5.76, 6.79] | 7.40E+03 | 3.89E+06 |
| SL-CoV | orf1a | 191 | 0.102<br>[0.0768, 0.126] | 0.460<br>[0.400, 0.516] | 2.04<br>[1.74, 2.34] | 4.45<br>[4.03, 4.84] | 3.00E+04 | 2.98E+07 |
| SL-CoV | orf1b | 191 | 0.0976 | n/a | n/a | n/a | n/a | 6.61E+00 |
| SL-CoV | S<br>protein | 191 | 0.132<br>[0.101, 0.161] | 0.561<br>[0.512, 0.605] | 1.13<br>[0.915, 1.37] | 2.02<br>[1.73, 2.35] | 3.00E+04 | 3.91E+07 |

### Correlated substitutions show evidence of recombination across specific genes in SARS-like betacoronaviruses

We used the 191 whole genome sequences used in the current *Nextstrain* build for SARS-like coronaviruses (SL-CoVs) (52–54) and aligned these to the NCBI reference genome for SARS-CoV-2 (see Materials and Methods for details). This included SARS-like coronaviruses from bats (BtCoVs), SARS-associated coronavirus or SARS-CoV-1 (SARS-CoV-1), and SARS-CoV-2 (SARS-CoV-2). We first examined whether there were hotspots of correlated substitutions along the length of the SL-CoV genome (Fig. 2A-B) in light of various reports of recombination hotspots in the SL-CoV and SARS-CoV-2 genomes, some of which suggest a correlation between adaptation and recombination in these regions (20, 22–25). We found correlated substitutions across the genome, with what visually appeared to be an accumulation of correlated substitutions in the coding sequence (CDS) region of orf1ab preceding the −1 ribosomal frameshift and the spike protein (throughout the manuscript, we will refer to the CDS region of orf1ab before the frameshift as “orf1a” and that after as “orf1b”).

**Figure 2.**
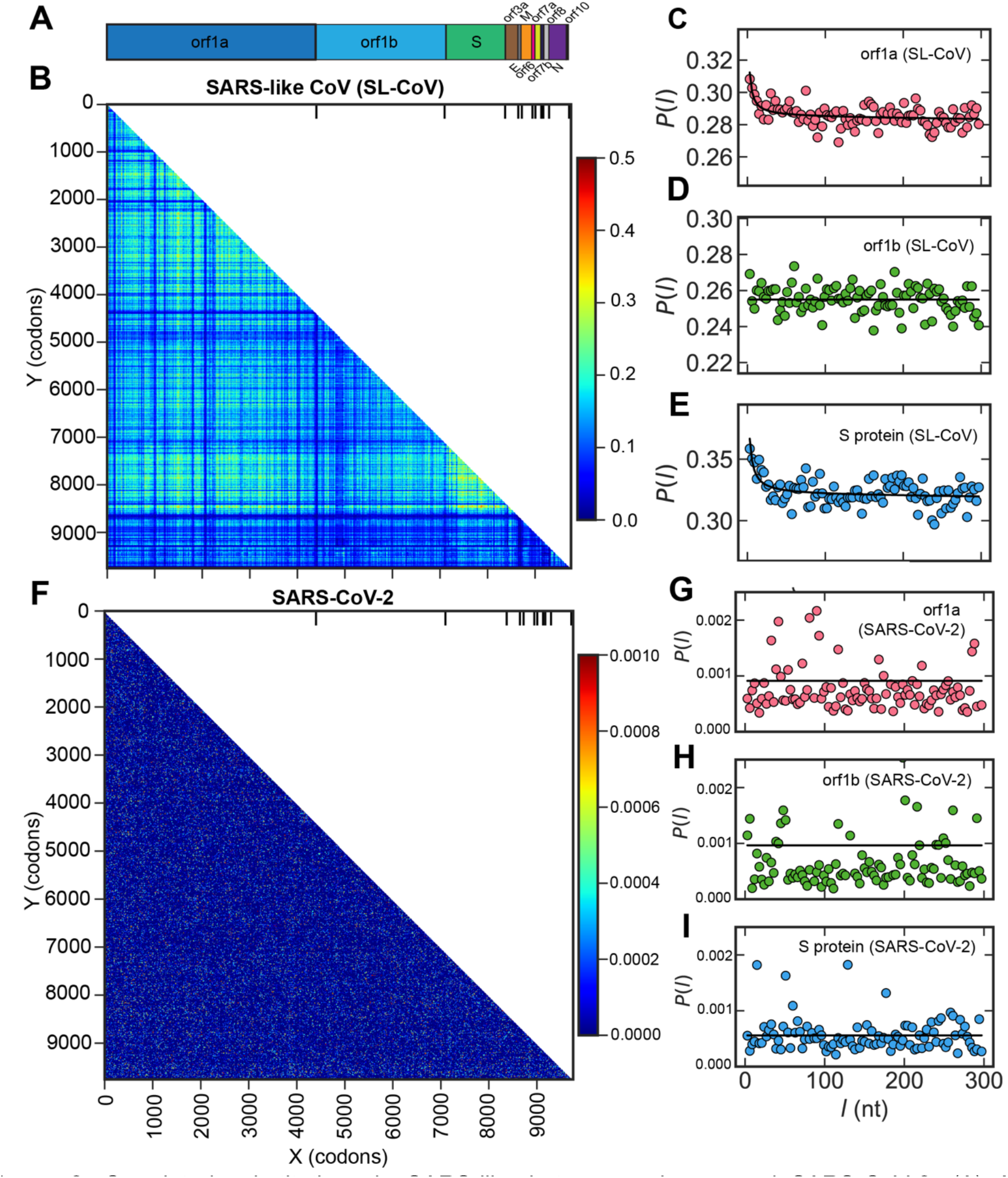
Correlated substitutions in SARS-like betacoronaviruses and SARS-CoV-2. (A) A schematic of the SARS-CoV-2 genome. (B) A heatmap of Pearson’s correlation coefficient (*ρ*(*X, Y*)) of synonymous substitutions across the coding regions of the SARS-like coronavirus genome constructed using 191 SARS-like betacoronavirus sequences (see Materials and Methods for more details). Each position (*X,Y*) displays the corresponding value of *ρ*(*X,Y*) as a color for a pair of codons located at genomic positions *X* and *Y* (given in codons). Coding regions are ordered genomically. Color bar indicates the value of *ρ*(*X,Y*). Ticks on the upper x-axis of the heatmap indicate where each CDS region begins and end, corresponding to the schematic of the SARS-CoV-2 genome above. “Orf1a” refers to the CDS region of orf1ab before the −1 ribosomal frameshift and “orf1b” refers to the CDS region after the frameshift. (C-E) Correlation profiles for the CDS regions of the orf1ab (C-D) and spike proteins (E) for SARS-like betacoronaviruses. Parameters of homologous recombination are given in Table 1. (F) A heatmap of *ρ*(*X, Y*) (analogous to *B*) but for 444,145 SARS-CoV-2 whole genome assemblies from NCBI (all available assemblies when the analysis was conducted). (G-I) Correlation profiles for the CDS regions of orf1ab (G-H) and spike proteins (I) for SARS-CoV-2. Inferred parameters are given in Table S4. In both heatmaps, monomorphic sites are assigned *ρ*(*X,Y*) = 0.

We next calculated correlation profiles and inferred recombination parameters across each gene (Fig. 2C-E, S3; Tables 1, S3) and found strong evidence for recombination in orf1a (Fig. 2C) and the spike protein (Fig. 2E). The CDS regions of the orf3a and N proteins also displayed evidence of recombination (Fig. S3), yet the shape of the decay in correlations was not nearly as apparent as those shown in Fig. 2. We observed orf1b had a similar synonymous diversity (*d_sample_*) to the ORF regions adjacent to it (Table S3), yet its correlation profile was flat (Fig. 2D); we hypothesize that the template-switching events occurring in the adjacent ORFs swapped out the entire orf1b CDS region, which would confer high diversity and a lack of recombination breakpoints. One CDS region which showed distinct patterns of correlated substitutions that cannot be adequately described by our model is that of orf8 (see Discussion). Overall, the inferred parameters suggest that the genes which show evidence of recombination are recombining frequently (Tables 1, S3); when considering (*γ/μ*)_*pool*_ inferred for the orf1a and S genes, the pools these samples have exchanged RNA with are rapidly recombining at rates ranging from ~2-5 recombination events per synonymous substitution.

Using every complete genome assembly for SARS-CoV-2 in the NCBI database (444,145 sequences at the time of this analysis), we measured correlated substitutions across the SARS-CoV-2 genome (Fig. 2F). In contrast to what we had observed in the SL-CoVs (Fig. 2B), we only detected very weak correlated substitutions across the SARS-CoV-2 genome. Correlation profiles across individual genes appeared to be largely flat (Fig. 2G-I, S4, inferred parameters in Table S4); this included the CDS regions for orf1 a and the spike protein (Fig. 2G,I), which showed signatures of recombination in the SL-CoV dataset (Fig. 2C,E). The pronounced difference in overall scale of *P*(*l*) between Fig. 2C-E and Fig. 2G-I reflects the differences in sample diversity between the SL-CoVs and SARS-CoV-2 (for these genes, *d_sample_* = 9.8 × 10^-2^ - 1.3 × 10^-1^ for the SL-CoVs and *d_sample_* = 6.0 × 10^-4^ - 1.2 × 10^-3^ for SARS-CoV-2). As new subvariants of SARS-CoV-2 arose during the peer review of this manuscript, we performed an updated analysis in July 2022 using *Nextstrain’s* subsampling of SARS-CoV-2 sequences from across the globe from the last six months (Fig. S5). The correlation profile measured across the genome of these sequences was still flat, with *d_sample_* = 1.2 × 10^-3^.

Recent work has suggested that SARS-CoV-2 experiences rate heterogeneity across the genome (27, 55), with specific genomic positions across the phylogeny exhibiting elevated mutation rates for *G* → *U* and *C* → *U* transitions, possibly related to APOBEC and ROS activity (55). This could cause individual sites to become ‘saturated’ (i.e., many identical mutations occurring at the same site across the tree) and specific genomic regions to exhibit anomalously high diversity, giving the appearance of recombination from a highly diverged source. If this effect were substantial, we would expect that the SARS-CoV-2 analysis presented in Figs. 2, S4, and S5 would show signatures of recombination, which they do not. To determine whether such effects impact our inference of recombination rates in other datasets, such as the SL-CoV dataset, we ran simulations using *phastSim* (56), which includes hypermutability models developed to simulate observed rate heterogeneity in SARS-CoV-2 (Fig. S6, details in Supplementary Information). In the simulations, we allowed a proportion of sites to be “hypermutable” and have highly elevated transition rates, with both the proportion of sites and the rates set to be equal to or exceed what has been estimated for SARS-CoV-2 (see Supplementary Information for details). We found that while sliding window averages of synonymous diversity increased in both magnitude and variability as expected (Fig. S6A-B), the correlation profiles we measured were consistently flat, correctly indicating that no recombination had occurred (Fig. S6C-D). These simulations suggest that heterogenous mutation rates, at least over a range which is biologically relevant to SARS-CoV-2, do not confound our ability to infer recombination rates using correlated synonymous substitutions.

### Clonal structure of the SARS-like betacoronaviruses

We sought to understand if a sufficient clonal signal remained in the SL-CoV samples which could be used to elucidate clonal relationships. We began by measuring genome-wide pairwise synonymous diversity (*d_sample_*) across the 191 SL-CoVs and clustering these sequences using the average linkage algorithm (57) to create a dendrogram (Fig. 3A; see Materials and Methods for details). We then split this tree into 11 flat clusters, where SARS-CoV-1 and SARS-CoV-2 each consisted of distinct clusters and the BtCoVs were broken into several clusters. The non-singleton BtCoV clusters were generally composed of sequences collected during the same time period and from the same geographic area; as examples, cluster 5 was almost entirely composed of samples from bats collected near Hong Kong and Gaungdong between 2005-2011 (58, 59), and cluster 6 was primarily samples collected near Yunnan Province from 2011-2014 (60, 61). As previously suggested (21, 25), it appears that on average across the genome SARS-CoV-2 is most closely related to a sequence cluster of BtCoVs (labeled “BtCoV (cluster 1)” in the legend shown in Fig. 3A). These two SL-CoVs were collected from bats in Zhejiang Province between 2015-2017 (62). Additional information pertaining to individual clusters is given with the sequence metadata (provided as a supplemental file).

**Figure 3.**
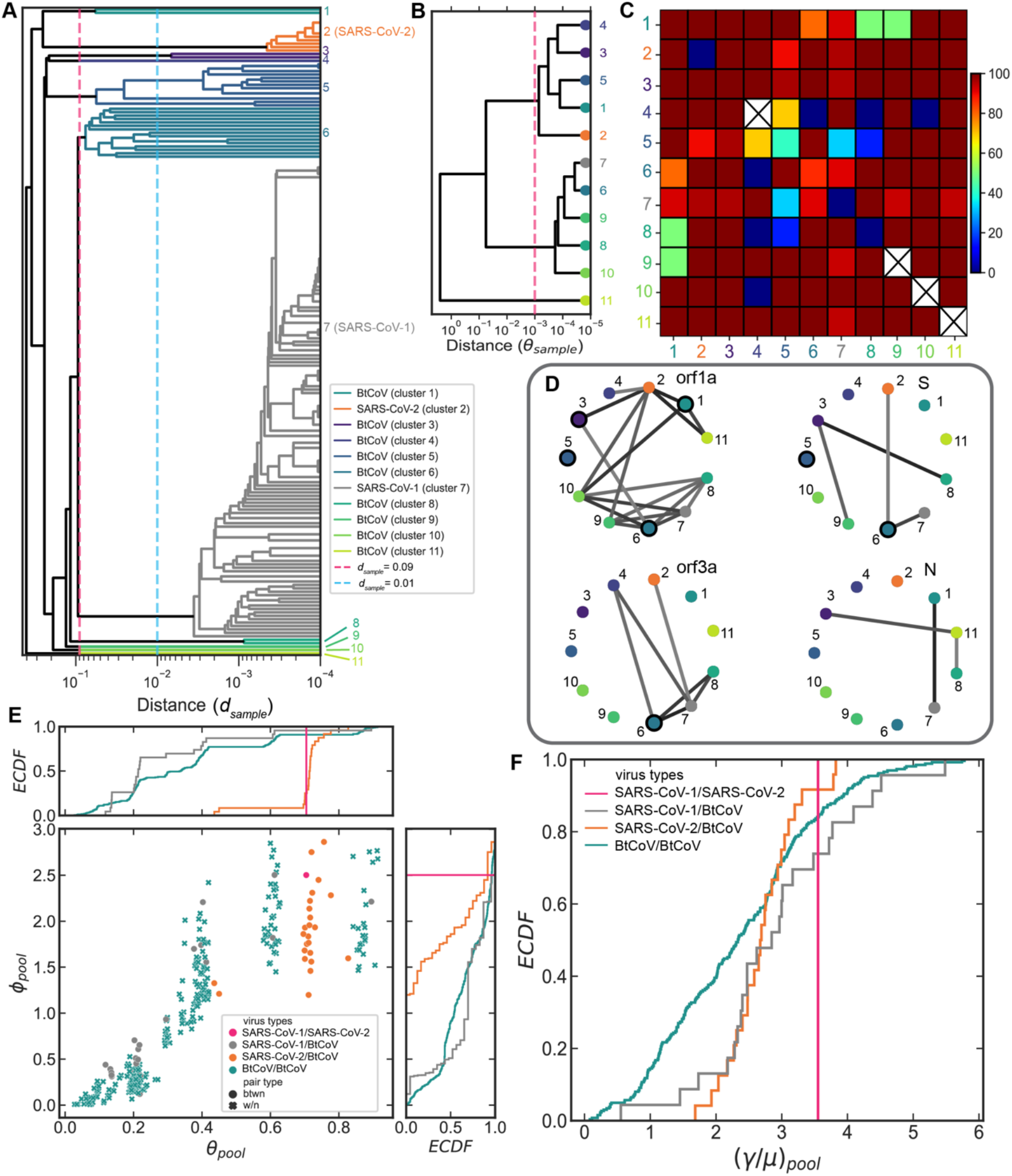
Pairwise analysis of recombination among SARS-like betacoronaviruses. (A) Dendrogram of the 191 SARS-like betacoronavirus sequences analyzed in both figs. 2-3. The tree was created using the average linkage algorithm with whole-genome pairwise synonymous diversity (*d_sample_*) as the distance metric (see Materials and Methods). The magenta, vertical dashed line depicts the distance at which the tree was cut to make the flat clusters shown in *B-C* (*d_sample_* = 0.09). The branch colors correspond to these clusters, as does the legend. The black, vertical dashed line depicts the cut at *d_sample_* = 0.01 made for the 27 flat clusters in *D-E*. Cluster numbers are shown along the vertical axis for the 11 flat clusters resulting from the cut made at *d_sample_* = 0.09. Horizontal axis is log-scale. Clusters composed of SARS-like coronaviruses from bats are labeled “BtCoV”. (B) Dendrogram of the 11 flat clusters from *A* created using the average linkage algorithm with *θ_sample_* as the distance metric. The red dashed line at *θ_sample_* = 10^-3^ indicates the maximum value beyond which the inference of *θ_sample_* is obscured due to recombination from the pool (as described in the Main text and Methods). (C) Heatmap depicting the percentage of total sequence pairs for a given pair of clusters which have recombined with a shared pool corresponding to the recombination network graph shown in Fig. S9. Sequence pairs were determined to have recombined with a shared pool by computing correlation profiles across the whole genome and fitting these profiles to the model. Diagonal depicts recombination between sequence pairs within the cluster. Clusters with single sequences have crosses through their diagonal cells. (D) Recombination networks for individual genes computed using correlation profiles calculated across each gene for pairs of clusters. Nodes are the 11 clusters from *A-B*, edges connect cluster pairs which have recombined with a shared pool. Black haloes around nodes indicate sequence pairs within the cluster have recombined. (E-F) Pool parameter distributions inferred from correlation profiles computed across the whole genome for pairs of sequence clusters. Clusters were made by cutting the dendrogram in *A* at *d_sample_* = 0.01 (depicted as vertical, black dashed line), resulting in 27 flat clusters. Distributions are separated by virus type; those distributions in which both clusters are within the same virus type are denoted as ‘w/n’, those which are between two virus types are denoted as ‘btwn’. In panel *D*, the main plot shows the pool recombinational divergence (*ϕ_pool_*) plotted against the pool mutational divergence (*θ_pool_*). Marginal plots show *ECDFs* of each pair’s divergence values. Panel *E* shows *ECDFs* of the relative recombination rate of the pool ((*γ/μ*)_*pool*_). All panels used the same fitting procedure as Figs. 1-2 (see Materials and Methods). For the recombination networks in *C*, if model selection suggested the profile was better fit with the null-recombination model, no edge was assigned for the cluster pair.

We then determined whether a statistically significant clonal signal remains in the sampled genomes by comparing the pool’s diversity (*d_pool_*), inferred from the correlation profile, to the sample’s diversity (*d_sample_*), which is measured from the sequencing data. In this case, we can use the mutational divergence (*θ_sample_*), which is proportional to the age of clonal portions, as a measure of clonal divergence. We computed the difference between *d_pool_* and *d_sample_* with respect to the variability in our measurement of *d_sample_*, a quantity which we refer to as the residual clonality (RC) effect size (see Eq. S3 and description in the Supplementary Information). For the 11 SL-CoV clusters, we first inferred recombination parameters for pairs of clusters (i.e., samples composed of sequence pairs in which neither sequence is from the same cluster). 51 out of 55 cluster pairs showed evidence of recombination as determined by model selection (see Methods; for the remaining 4 cluster pairs, *θ_sample_* was determined via standard population genetic expression for heterozygosity (63) given by Eq. 5). We then plotted the recombination coverage and *θ_sample_* for these cluster pairs against the RC effect size (Fig. S7), and determined that when *θ_sample_* was greater than ~10 ^3^, the RC effect size was generally < 1. Therefore, for this dataset, we are able to infer values of *θ_sample_* < 10^-3^, while for cluster pairs having RC < 1 we can confidently conclude only that *θ_sample_* > 10^-3^.

We constructed an average linkage tree based on *θ_sample_* for the 11 SL-CoV clusters, and demarcated the value above which *θ_sample_* is not well-determined (Fig. 3B). We found that there is sufficient residual clonality in the data to infer the clonal structure for most of the SL-CoV lineages (Fig. 3B), revealing key differences with respect to the dendrogram based on genome-wide pairwise distances (Fig. 3A). While the tree in Fig. 3A suggests that SARS-CoV-2 shares its most recent common ancestor (MRCA) with the BtCoVs of cluster 1, the clonal tree in Fig. 3B indicates that SARS-CoV-2 actually shares a most recent common ancestor (MRCA) with clusters 1, 3, 4, and 5. Furthermore, this means that SARS-CoV-1 does not share its MRCA with clusters 3-5 in the clonal tree.

To determine whether standard phylogenetic methods which assume no homologous recombination recover this clonal structure, we used *IQ-Tree* (64), which relies on maximum likelihood estimation for phylogenetic inference, to reconstruct the phylogeny for the sequences depicted in Fig. 3A using a generalized time reversible (GTR) model (Fig. S8; see Materials and Methods for details). We found that the structure of the tree matched Fig. 3A, but not that of Fig. 3B. This may suggest that, for the SL-CoVs, standard phylogenetic inference yields phylogenies which are likely obscured by recombination; the resultant phylogenies reflect both recombination from the pool and mutations within the sample.

### Correlated substitutions reveal the gene pool structure of SARS-like betacoronaviruses

To determine which members of the SL-CoVs have recombined with shared gene pools, we calculated correlation profiles across the whole genome for all possible sequence pairs and fit each profile to the model. We then visualized recombination between sequence pairs as a network graph (Fig. S9), where each strain is a node, and edges connect strains which have recombined with a shared pool. Visually, the network appears to be highly connected, suggesting that many pairs have recombined with shared gene pools. To quantify network connectivity, we created a matrix of the percentage of sequence pairs which have recombined with a shared pool for a given cluster pair (Fig. 3C). This analysis reveals that the clusters are less interconnected than they appear, as not all clusters share pools equally; and in some cases, we find clusters that may not share the same gene pool at all. This indicates that distinct, structured gene pools exist despite a high degree of recombination and gene pool sharing across the SL-CoV lineages.

Because our analysis of the SL-CoVs revealed that individual genes have different recombination parameters, we tested whether the four genes which showed recombination signatures in Fig. S3 each had distinct networks of recombination events. For each of the four genes, we measured correlation profiles for pairs of sequence clusters and used these profiles to determine if the cluster pair showed evidence of recombination with a shared pool (Fig. 3D). As profiles measured over single genes account for fewer genomic sites compared to profiles measured over whole genomes, the recombination signal for individual genes needs to be stronger relative to random correlations to allow for detection; this accounts for the slight discrepancies between Fig. 3C and D. Each of the genes had a unique recombination network, with the most recombination occurring in orf1a. Furthermore, the analysis suggested SARS-CoV-1 and SARS-CoV-2 both underwent recombination events with the same pool in the orf3a CDS region.

To further examine the structure of the SL-CoV gene pools, we cut the tree in Fig. 3A at *d_s_* = 0.01 (yielding 27 clusters) and computed correlation profiles for each cluster and cluster pair across the entire genome and inferred their recombination parameters. We first investigated the degree of clonality of each sample (a cluster or cluster pair) by computing its RC effect size (see Methods and previous section). As we have a larger sample distribution (i.e., more cluster pairs) than in the previous section, we can adopt an even stricter criterion for the RC effect size here; if we specify that the RC effect size must be greater than 2 to infer *θ_sample_*, by plotting *θ_sample_* versus the RC effect size we determined that 82% of samples had a sufficient RC effect size (Fig. S10). For these samples, *θ_sample_* ranged from 1.8e-5 to 3.8e-3 (we note the upper bound is similar to that used in the previous section), with a median of 1.6e-4, while *c_sample_* ranged from 27% to 97% with a median of 87% (Fig. S10). For the remaining 18% of samples where RC effect size ≤ 2, recombination coverage was comparatively higher, with *c_sample_* ranging from 92% to 100%, and a median of 98%, and for these samples we estimate a lower bound of *θ_sample_* ~ 3.6e-4. Across all SL-CoV samples, we find the median *θ_sample_* = 1.9e-4 and median *c_sample_* = 92%. To test that the inferred parameters provide reasonable estimates for *θ_sample_* and *c_sample_*, we took several sequence pairs with varying levels of *c_sample_* and looked at sliding window averages of diversity 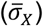 across the genome (Fig. S11). Our estimates of *θ_sample_* suggest that clonal regions should exhibit diversity levels in the range ~1e-3 to 1e-5, and we found that the genomic fraction with 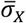 in this range roughly matched the inferred clonal fraction, 1 – *c_sample_*. The remainder of the genome has been recombined in from the pool, and we find that the average diversity of these regions is close to our estimate of the pool diversity. Moreover, we can use the distribution of zero-SNP block lengths in these sequence pairs to provide an alternative estimate of *c_sample_*, and find that this roughly matches the values inferred from the coalescent model (Fig. S12, see Supplementary Information for details). We conclude that despite the extensive recombination across the SL-CoV lineages, a substantial clonal signal remains in the data and a wide distribution of clonal divergence times, spanning at least two orders of magnitude, can be detected.

We then sorted each of the samples according to its virus type (e.g., a cluster pair in which one cluster is comprised of BtCoV sequences and the other is SARS-COV-1 sequences is labeled “SARS-CoV-1/BtCoV”), and examined the distributions of *ϕ_pool_* and *θ_pool_* across samples. We found that each pair of virus types had a distinct parameter distribution (Fig. 3E). Moreover, the divergence distributions are multi-modal (particularly the BtCoV/BtCoV distribution), suggesting that subsets of these samples interact with gene pools with distinct evolutionary dynamics. To remove the dependence on coalescence time, we plotted the distributions of (*γ/μ*)_*pool*_ for each cluster pair as empirical cumulative distribution functions to assess the relative recombination rates of these pools (Fig. 3F). This revealed that within the BtCoV lineages’ pools, there is a wide and relatively uniform distribution of recombination rates, while each of the SARS-CoV-1 and SARS-CoV-2 recombine with shared BtCoV pools with similar characteristic rates, and exhibit narrower, unimodal distributions. Nearly all cluster pairs have (*γ/μ*)_*pool*_ > 1, meaning that in SL-CoV gene pools there are multiple recombination events occurring per synonymous substitution, indicating that recombination plays a major role in the evolution of SARS-like coronaviruses.

## Discussion

We adapted the non-phylogenetic, computationally-efficient *mcorr* method (originally developed for analysis of bacterial genomes (29, 30)) to infer the parameters of homologous recombination for SARS-like coronaviruses and other RNA viruses. The methodological advances reported here include the use of a two-parameter template-switching model, the introduction of the residual clonality effect size as a tool for clonal inference, the ability to analyze single genes, the measurement of correlation profiles across whole genomes (vs gene-averaged profiles), and the improved efficiency of the method such that >400,000 sequences can be analyzed. We first demonstrated that this method is generally applicable to (+)ssRNA viruses by inferring recombination parameters for viruses which have known histories of recombination using datasets consisting of hundreds to thousands of WGS. We then applied this to understanding recombination in SARS-like coronaviruses. We found strong signatures of recombination in the CDS regions of orf1a and the spike protein. While previous studies of SARS-like coronaviruses have yielded estimates of recombination rates among analyzed sample sequences (20) and others have suggested that SARS-like coronaviruses have recombined with unsampled pools (21, 24), here we infer recombination rates and parameters for both the sample sequences and the unsampled gene pools with which they recombine.

Our gene-by-gene analysis of recombination for the SARS-like coronaviruses revealed that orf1ab and the S protein show strong signatures of recombination, and suggested that these parts of the genome recombine at high rates, ranging from ~2-5 recombination events per synonymous substitution (Table 1). Interestingly, when we fit the correlation profile for orf1a with the fragment-incorporation model, we found that the 95% bootstrap confidence interval for the mean fragment size ranges from ~11,000 to 30,000 nt, suggesting that the SL-CoVs may take up fragments via recombination, consistent with previous experimental observations with betacoronaviruses (34) (for additional discussion see section on zero-SNP blocks in Supplementary Information). We note that orf8 showed distinct patterns of correlated substitutions that cannot be adequately described by our model (Fig. S3). This region is thought to be highly variable, to contain several stem-loops which could lead to correlations between distant sites, and to have undergone recombination (65–67). Furthermore, the RNA secondary structure in this region could lead to selection on synonymous sites resulting in codon bias (68). Therefore, we speculate that this combination of RNA secondary structure and recombination has left the orf8 gene with an uncharacteristic decay in correlated substitutions; additionally, the decay we observe could be impacted by the many deletions and nonsense substitutions in this region (66, 67). Because we cannot reliably infer parameters using our recombination model for this CDS region, and we also cannot assume that evolution has proceeded clonally in this region, for orf8 we simply list *d_sample_* for this gene (given in Table S3).

By measuring correlated substitutions between SL-CoV sequence clusters and decoupling the contributions of mutation and recombination to the sample diversity, we were able to recover much of the clonal structure for the analyzed SL-CoV clusters (Fig. 3B). We show that due to both high recombination coverage and variability in the measurement of *d_sample_*, the residual clonal signal in the data only allows sample ages up to *θ_sample_* ~ 0.001 to be determined. Nevertheless, this is sufficient to yield important insights into the clonal relationships between the sequence clusters. Our inference of clonal relationships indicates that SARS-CoV-2 shared an MRCA with BtCoVs from clusters 1, 3, 4, and 5, whereas clustering based on *d_sample_* (Fig. 3A) and standard methods for phylogenetic inference such as maximum likelihood estimation (Fig. S8) would suggest that SARS-CoV-2 shares its MRCA with cluster 1 only. Whereas phylogenetic methods attempt to identify ancestral relations by inferring or simulating individual recombination and mutation events that took place in different portions of the genome at different times in the past (see below), our approach determines the residual clonality of a pair of clusters directly from its correlation profile. In this respect, our method of clonal inference is more direct, however future analyses using broader sets of SL-CoV sequences, in combination with simulations and experiments, will be needed to determine when each approach may be most advantageous. By measuring correlated substitutions between individual sequence pairs, we inferred a recombination network for the SL-CoVs and found that the majority of sequences have recombined with a pool during their evolutionary history (Fig. S9). Counting the number of recombined pairs (Fig. 3C) shows that some clusters appear to have only recombined with subsets of sequences from other clusters (e.g., cluster 5), indicating that while the SL-CoV recombination network is dense, it is also heterogeneous. We further found that each gene had a unique recombination network (Fig. 3D) suggesting each region has been shaped by different sets of recombination events along the evolutionary trajectories of the samples.

The observed heterogeneity in the connectivity of sequence clusters in these recombination networks led us to hypothesize that the gene pools which SL-CoVs recombine with are partitioned or structured. We tested this hypothesis by inferring recombination parameters of the SL-CoV gene pools and found these to be diverse and characterized by high recombination rates (Fig. 3E-F). Our inference of the corresponding *θ_sample_* distribution (Fig. S10B) shows that generally *θ_pool_* is orders of magnitude higher, which suggests that the SL-CoV gene pools constitute a diverse and largely unsampled reservoir of viral sequences. Moreover, the *c_sample_* distribution (Fig. S10A) indicates that the set of SL-CoV genomes have been substantially impacted by recombination, with a median of *c_sample_* ~ 87% (for RC effect size > 2). The diversity and structure of the gene pools for a given microorganism can vary widely; we can imagine a scenario in which different samples from a microbial population interact with a discrete set of gene pools or, alternatively, these gene pools could overlap, leading to recombination events occurring between a sample and multiple pools (Fig. 4A). What sets the ‘softness’ of gene pool boundaries in microorganisms is unclear, and will undoubtably be the focus of future investigations. In the case of SL-CoVs, it seems unlikely, based on the literature (24, 25, 69, 70), that the molecular mechanisms underlying recombination and mutation are so unique to each member of this group of viruses that it would result in the diverse distributions of pool parameters which we observe (Fig. 3E-F). However, SL-CoVs can exist in a broad range of hosts with different sets of selective pressures (11, 70), and these diverse environments with unique selective pressures may strongly influence the observed recombination rates.

**Figure 4.**
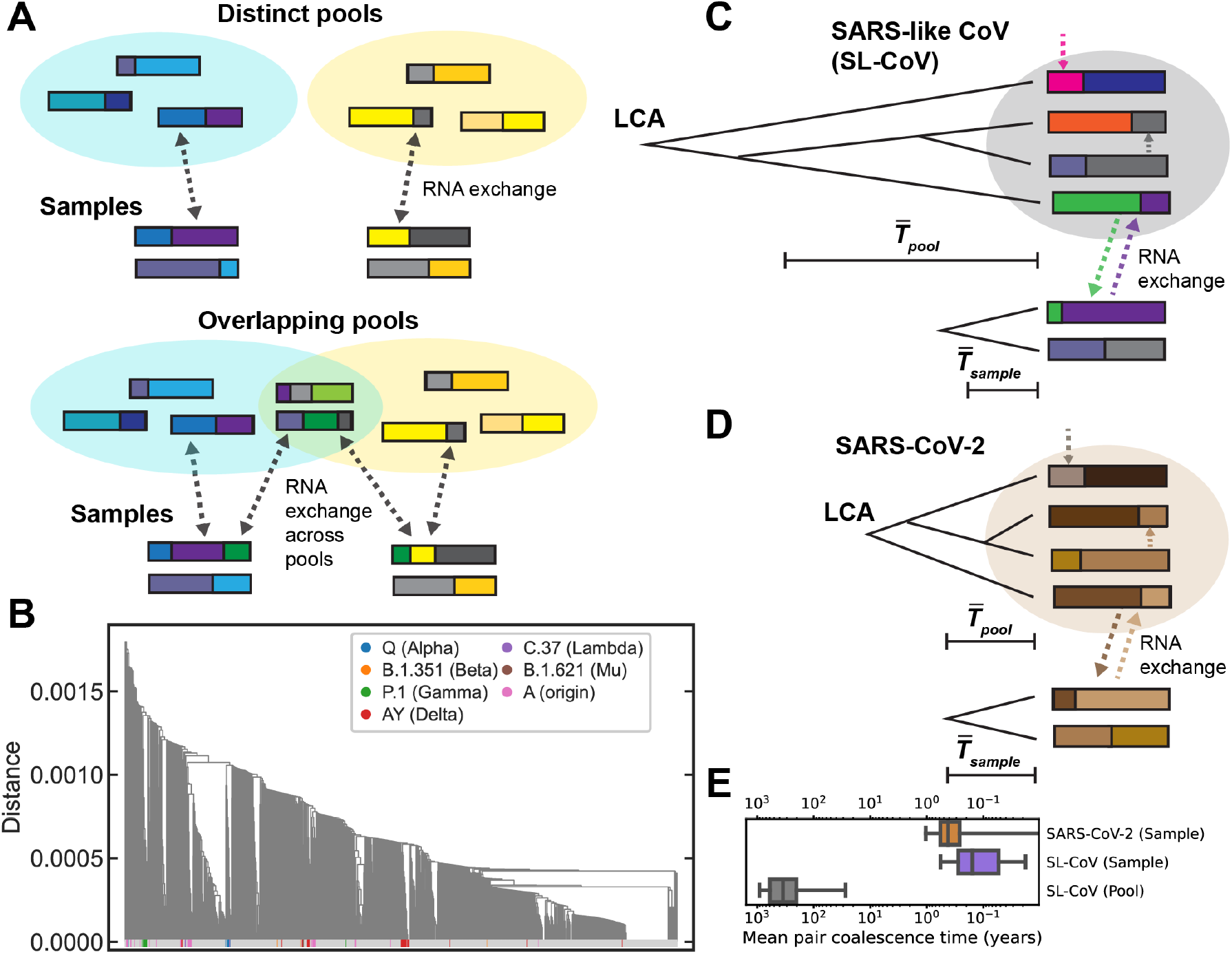
Gene pool structure and masking of recombination by sequence homology. (A) Schematic depicting samples recombining with distinct gene pools which have no overlap, and overlapping gene pools where recombination can occur between multiple pools and a given sample. (B) Dendrogram of SARS-CoV-2 sequences from the NCBI database, where one sequence from each Pango lineage was randomly selected to represent that lineage. Pairwise distances were computed using genome-wide synonymous diversity (see Materials and Methods) and clustering was performed with the average linkage algorithm. Lineages whose parent lineages are World Health Organization designated Variants of Concern and Interest (as of October 27, 2021 (92)) have colored tips, all other tips are colored gray. (C-D) Schematics illustrating hypothesis for why detecting recombination using correlated substitutions is not possible using just SARS-CoV-2 sequences. Panel *C* shows SARS-like coronavirus (SL-CoV) samples recombining with a diverse pool, and panel *D* shows SARS-CoV-2 recombining with a pool of SARS-CoV-2 sequences. LCA is last common ancestor. (E) Tukey boxplots of the distributions of coalescence times of sequence pairs for samples and pools corresponding to the schematics in *C-D*. For the SARS-like coronaviruses (SL-CoV), the distributions are the mean coalescence times for sequence pairs within each of the clusters and cluster pairs shown in Fig. 3E. For the calculation of 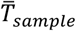 of the SL-CoVs, only clusters and cluster pairs with RC effect size > 2 were used (see Main text and Eq. S3 of Supplementary Information). For SARS-CoV-2, the boxplot depicts the distribution of coalescence times for the sequence pairs of the tree shown in Fig. 4B. The line bisecting the box is the 50^th^ percentile, the upper and lower edges of the box are the 25^th^ and 75^th^ percentile respectively, and the whiskers are 1.5*IQR. Horizontal axis is logarithmic (the lower whisker for SARS-CoV-2 extends to zero, which is not shown).

A potential confounding factor in our analysis is heterogeneous mutation rates, as it has been demonstrated that SARS-CoV-2 experiences rate heterogeneity across the genome (27, 55). By analyzing single genes (Figs. 2C-E and S3), we account for variability in mutation rates across large genomic regions. By running simulations with heterogenous mutation rates and hypermutable sites, we control for finer scale heterogeneity in mutation rate, and our simulations suggest that, at least over a biologically relevant range, these effects do not confound our analysis (Fig. S6). Additionally, it has been shown that many human pathogenic RNA viruses exhibit heterogeneous coalescence times as a consequence of variation in selection over time (71). We have controlled for this by separately inferring pool recombination rates for individual SL-CoV sequence clusters or cluster pairs (Fig. 3F), for which coalescent times are much more tightly distributed, or across the entire sample phylogeny for which coalescence times are heterogenous (tree in Fig. 3A, recombination rates in Table 1 for orf1a and the spike protein); the inferred pool recombination rates are similar, indicating that variability in coalescence time does not substantially impact our inference of recombination rates for gene pools. It is possible that selection on synonymous sites, e.g., relating to codon usage bias and preferences for CpG dinucleotide frequency and GC richness, could affect our analysis (55, 72–75). However, the strength of such selection in SL-CoV is unclear; recent work found primarily statistically insignificant patterns with regards to selection relating to CpG and GC content (55), and there have been results suggesting there is selection both against (76) or for (55) U content. In previous work we ran simulations with an analogous model which showed that selection at linked sites, which can act to reduce diversity at synonymous sites, minimally affects our analysis (29).

Our analysis of the SL-CoV samples indicates that recombination occurs at least as often as mutation in nearly all lineages [(*γ/μ*)_*pool*_ > 1; see Fig. 3F], a result that differs substantially from inference based on Bayesian MCMC phylogenetic simulations on a similar SL-CoV dataset, which found that recombination events occur 200 times less frequently than mutations (20). Phylogenetic methods typically attempt to infer the ancestry of each piece of DNA within the sampled genomes by modeling all possible recombination and mutation events that could have occurred since its coalescence. Inference of the maximum likelihood set of recombination events relies on the existence of inconsistencies in the inferred tree of different pieces of DNA across the sample. As branches are joined going backward in time, the size of the trees decreases monotonically, and there is progressively less evidence to call recombination events. Such methods are thus expected to underestimate recombination rates in strongly clustered samples with very deep branches, such as the SL-CoV dataset (Fig. 3A). In contrast, our approach accounts for recombination events that occur in the external, unsampled pool; these events cannot be individually inferred, however their signature is the correlation profile. These correlations develop over long timescales under the combined effect of historical recombination and mutation events in the pool (29), and may enable more accurate measurements of (*γ/μ*)_*pool*_ for deeply branched samples (30). Additionally, the recombination rate estimated in ref. (20) is limited to recombination during co-infection events between distinct viral lineages, while the pool recombination rate we estimate here is based on all recombination events that occur in the ancestry of the viral gene pool, including within-host recombination.

We analyzed every complete genome assembly for SARS-CoV-2 from human hosts in NCBI (444,145 at the time of analysis) and created a map of correlated substitutions across the genome (Fig. 2F). We did not observe any regions with strongly correlated substitutions, nor did we find that any genes had correlation profiles which indicated the presence of recombination (Fig. 2G-I, S4). At first this may give the impression that SARS-CoV-2 has recombined little since it entered the human population in late 2019. However, given (i) the high recombination rates of related SL-CoV strains, (ii) the high levels of ancestral recombination we measured between SARS-CoV-2 and SL-CoV, and (iii) the conserved molecular mechanism of RNA replication which underlies template switching, we hypothesize that recombination is most likely occurring among SARS-CoV-2 in human hosts yet insufficient time has passed for SARS-CoV-2 to accumulate enough diversity to allow for detection of recombination via correlated substitutions. A simple comparison of an average linkage dendrogram sampling across all major lineages of SARS-CoV-2 (Fig. 4B) shows that the overall diversity levels are orders of magnitude lower for SARS-CoV-2 as compared to the SARS-like coronavirus dendrogram (Fig. 3A). Other studies have previously suggested that the current lack of SARS-CoV-2 diversity impedes the ability to detect recombination (77, 78) resulting in what these investigators suggest are potentially large underestimates of recombination levels in these pandemic datasets (79).

We can further this argument by estimating differences in coalescence times for SARS-CoV-2 versus the SARS-like coronaviruses as a whole. We use a standard maximum likelihood phylodynamic approach (*TreeTime;* (54)) to estimate the mutation rate as μ≈ 9.8 × 10^-4^ *bp*^-1^ · *year*^-1^ for SARS-CoV-2 (see Methods for details); for the SARS-like coronaviruses, previous studies have inferred μ ≈ 5.0 × 10^-4^ *bp*^-1^ · *year*^-1^ (20, 21). We use these mutation rates along with *θ_sample_* and *θ_pool_* to estimate the mean coalescence times for pairs in the sample and pool for the SL-CoVs and SARS-CoV-2. For the SL-CoVs, if we use the median values of *θ_sample_* and *θ_pool_* for the parameter distributions of the 27 clusters appearing in Fig. 3E-F, we find that 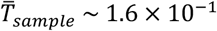 years and 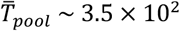 years (Fig. 4E; for 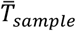, we use cluster pairs for which RC effect size > 2, as described in Results). This suggests that the SL-CoV samples recombined with pools which have been accumulating diversity for much longer times than the samples (Fig. 4C). For SARS-CoV-2, we can use the *θ_sample_* distribution of the SARS-CoV-2 sequence pairs in the dendrogram in Fig. 4B to compute the median coalescence time for pairs as: 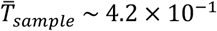 years (Fig. 4E; *θ_sample_* was computed for each sequence pair using the classic population genetics expression for pairwise heterozygosity (63) given as Eq. 5 in Methods). For this dataset, which is comprised solely of SARS-CoV-2 sequences from human hosts, opportunities for recombination have almost exclusively stemmed from co-infection events involving multiple strains from local transmission chains between humans, in which every RNA sequence is highly similar (Fig. 4D). Furthermore, in the case of this SARS-CoV-2 dataset, we know that our sample has effectively the same coalescence time and rate of synonymous substitution as the pool, because sequences in the sample and pool are highly overlapping (the pool here being un-sequenced SARS-CoV-2 strains in the human population). If we therefore use 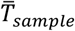 as our estimate of 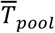 in SARS-CoV-2, this suggests that, consistent with expectation, the SL-CoV pools have had much longer to accumulate diversity, which allows us to differentiate between two sequences which have swapped to create a new hybrid when analyzing correlated substitutions. While we cannot predict when sufficient diversity will have accumulated to allow for the detection of recombination via correlated substitutions, 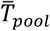 for the SL-CoVs suggests this may be on the order of ~10^2^ years. This analysis therefore further emphasizes the current need for computationally-efficient approaches to sift through the massive amounts of available sequencing data to pinpoint recombinant SARS-CoV-2 sequences (e.g., (78, 80–82)).

While previous studies have examined recombination in SARS-like coronaviruses via the analysis of phylogenetic incongruence and Bayesian inference (20–25), our work here offers unique advantages and insights; it makes no assumptions of phylogenetic structure or evolutionary parameters (i.e., specification of prior distributions for parameters which is necessary for Bayesian inference), and we infer population genetic parameters of the larger, unsampled viral reservoirs that the SARS-like coronaviruses have recombined with. Moreover, we use differences in diversity levels between a sample and its pool to determine the residual clonal signal in the data. Our methodology enables analysis of the massive datasets that have become the norm in COVID-19 epidemiology, and which are prohibitively large for current Bayesian simulation-based approaches. Our work yields a new set of tools to analyze recombination in positive-sense RNA viruses, and reveals the parameters of homologous recombination of the diverse set of gene pools with which SARS-like coronaviruses recombine. This may aid in understanding how the interplay among population structure, selection, and recombination acts to mold the unique genetic architecture of viruses at the center of major epidemics.

## Materials and Methods

### Data and code availability

For complete genome assemblies from NCBI that were used in this manuscript, we provide lists of Genbank accession numbers as supplemental files to this manuscript. These genome assemblies can be retrieved by uploading the accession lists to NCBI Batch Entrez (https://www.ncbi.nlm.nih.gov/sites/batchentrez) which will locate the sequences in the NCBI database. For the 191 SARS-like coronavirus sequences used in figs. 2-3, we provide both the original unaligned, multi-fasta file from the *Nextstrain build* (the GitHub repository for this build can be found here: https://github.com/blab/sars-like-cov) and the aligned extended multi-fasta (XMFA) file and the corresponding metadata for the sequences in the event that this GitHub repository becomes inactive. We removed the strain labeled “Modified_Microbial_Nucleic_Acid” as it was unclear was modified in the laboratory. In the XMFA file “APS229_sl-cov_leq10_gaps_placeholders.xmfa”, any gene sequence with <90% alignment to the SARS-CoV-2 reference genome has been replaced with dashes as placeholders (necessary for the scripts *mcorrLDGenome, mcorrPairGenome*, and *calcKsPair*). The XMFA file “APS229_sl-cov_leq10_gaps_no_placeholder.xmfa” simply removes any gene sequence with <90% alignment (preferable if using *mcorrGeneAln* and performing bootstrapping). In the metadata file, we included the cluster number for each sequence for the two different cuts of the dendrogram in Fig. 3A. Those sequences which do not have cluster numbers are not publicly available through GenBank, and were not included in the analysis. For the dendrogram appearing in Fig. 4A, we provide an XMFA file for the SARS-CoV-2 genomes used to construct it. Genbank accession numbers for all reference genomes used in this study are listed in Table S5. Results from fitting correlation profiles of individual SL-CoV sequence pairs and the sets of 11 and 27 SL-CoV sequence clusters from Fig. 3 and 4 are provided as supplemental .csv files. For the SARS-CoV-2 analysis using the *Nextstrain* build as of July 13^th^, 2022 appearing in Fig. S5 (we used the global analysis built with GenBank data), we provide the aligned XMFA file and corresponding metadata for the sequences. All original code for this study has been deposited in GitHub and will be made publicly available as of the date of publication. Unless otherwise specified in the *Materials and Methods* section of the manuscript, all original code for the study can be found here: https://github.com/kussell-lab/viral-mcorr.

### Generation of multi-sequence alignment files

For all RNA viruses studied, we used reference-guided alignment to build consensus genomes by taking whole genome assemblies and aligning them to a reference genome from NCBI (Genbank accessions for reference genomes listed in Table S5) using the program *ViralMSA* (83) with *Minimap2* as the aligner (84). We then used our in-house program *splitFasta* to split the multi-FASTA file generated by *ViralMSA* into separate FASTA files for each genome, and used our program *CollectGeneAlignments* to extract CDS regions and generate an XMFA file. We filtered out any gene alignment with >10% gaps using our program *FilterGaps*. For calculations of correlation profiles across single genes, our program *geneMSA* was used to split the XMFA file including all gene CDS regions into separate multi-fasta files for each gene, which could then be analyzed with our program *mcorr-gene-aln* (described below). We created the program *CollectGeneAlignments* previously (used in (30, 31) and it can be found here: https://github.com/kussell-lab/ReferenceAlignmentGenerator. All other in-house programs can be found here: https://github.com/kussell-lab/viral-mcorr.

### Measurement of correlation coefficient of synonymous substitutions

We computed Pearson’s correlation coefficient for synonymous substitutions along the length of the genome using the following expression:

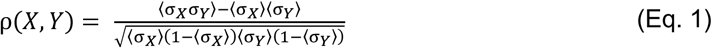

where ρ(*X,Y*) is Pearson’s correlation coefficient for a pair of codons at genomic positions *X* and *Y*, and *σ_i_* is the substitution profile for a site *i* (as described in the main text, this is a binary variable assigned a 1 for difference and 0 for identity at genomic position i). The substitution profile is measured at each four-fold degenerate, third-position codon for every sequence pair fc, then averaging over all pairs. When performing this calculation, if for a given sequence pair either site *i* = *X* or *Y* had a gap, this sequence pair was not included in the calculation of 〈σ_*X*_〉, 〈σ_*Y*_〉, or 〈σ_*X*_σ_*Y*_〉. This is because the discrepancy could result in different sets of sequence pairs being used in the calculation of these mean substitution profiles, which in some cases allows ρ(*X,Y*) to range outside of −1 < ρ < 1. For the *Poliovirus* dataset (Fig. 1C) and SARS-like coronavirus dataset (Fig. 2B), we developed the program *mcorrLDGenome* to perform the calculation of these substitution profiles. Because the calculation of ρ(*X, Y*) for the SARS-CoV-2 heatmap in Fig. 2F is very memory-intensive given the number of sequences, we created a program called *makeGeneDB*, which takes the aligned sequences (in XMFA format) and stores them in local key/value databases. These can then be accessed by the program we developed called *mcorrLDGenomeLite*, to perform the calculation in a memory-light manner. This allows for a short, memory-intensive calculation performed by *makeGeneDB*, followed by a longer, memory-light calculation performed by *mcorrLDGenomeLite*. To perform the calculations of ρ(*X, Y*) for the coronaviruses (both the SARS-like coronaviruses and SARS-CoV-2), when we filtered out gene alignments with >10% gaps, *FilterGaps* was used with the flag “fill-gaps” on; this is a requirement for both *mcorrLDGenome* and *mcorrLDGenomeLite*, which sort the CDS regions by the position they appear in the genome into one continuous sequence of codons. Additionally, when *CollectGeneAlignments* is used for the coronaviruses, the *orf1ab* CDS region appears three separate times in the NCBI reference genome’s GFF3 file: 266 to 13468 (CDS region for pp1ab), 13468 to 21555 (continuation of CDS region for pp1ab), and 266 to 13483 (CDS region for pp1a). This results in overlapping CDS regions in the resultant XMFA file. To create one, genomically-ordered CDS region, we removed the CDS region for pp1a from the XMFA files generated by *CollectGeneAlignments* (which is for our purposes redundant to the CDS region for pp1ab) before performing calculations with *mcorrLDGenome* or *mcorrLDGenomeLite*. For simplicity, in all figures in the manuscript, the CDS region from 266 to 13468 is labeled “orf1a” and the region from 13468-21555 is labeled “orf1b”. To facilitate reproducibility, we have also provided an example Jupyter Notebook which shows how to use the output of *mcorrLDGenome* and *mcorrLDGenomeLite* to create the heatmap shown in Fig. 1C (in the *viral-mcorr* github repository see: “pcc_example.ipynb”).

### Measurement of sample correlation profiles for single genes and whole genomes

Using the whole genome alignments of our sample sequences we measure the ‘substitution profile’ (*σ_i_*(*k*)) at each four-fold degenerate, third-codon position *i* for every sequence pair *k*. When calculating correlation profiles for single genes, we do this separately for each gene’s CDS region, where all viruses appearing in this manuscript except for the coronaviruses only code for a single polyprotein. When computing correlation profiles across the genome of the coronaviruses (e.g., Fig. S9, 3D, and 3E), we first sort the CDS regions by the position they appear in the genome into one continuous sequence of codons, and then compute correlation profiles across the entire genome. We compute the pairwise synonymous diversity of a CDS region as:

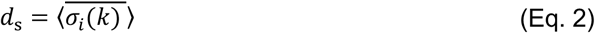

in which the bar signifies averaging over sequence pairs *k* and the bracket signifies averaging over positions *i*. We compute the joint probability of synonymous substitutions for a pair of sites separated by *l* nucleotides as:

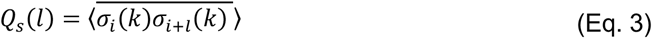

The correlation profile is then calculated as *P*(*l*) = *Q_s_*(*l*)/*d_s_*. For single genes, these computations were performed using the command-line (CLI) program *mcorr-gene-aln* (developed for this manuscript), which takes as input an alignment for a single CDS region formatted as an extended multi-fasta (XMFA) file. Bootstrap replicates of the correlation profile were created by sampling the alleles for a gene with replacement. For the computation of correlation profiles for SARS-CoV-2 genes shown in Fig. 2 and S3, *mcorr-gene-aln* was used with the number of bootstraps set to zero, with the exception of “orf1a” (Fig. 2G and top-left panel of Fig. S3). For the calculation of this correlation profile, using *mcorr-gene-aln* was highly memory-intensive given the number of sequences and sequence length; therefore, we developed the program *mcorr-gene-lite* to access the aligned sequences via local key/value databases created by *makeGeneDB* (described in “Measurement of correlation coefficient of synonymous substitutions”) and perform the calculation in a memory-efficient manner. For the computations of correlation profiles across the whole SARS-like coronavirus genome for individual sequence pairs in Fig. S9, the program *mcorrPairGenome* was developed. For computations of correlation profiles across the whole SARS-like coronavirus genome for sequence clusters and pairs of sequence clusters, the program *mcorrViralGenome* was developed. When calculating correlation profiles between pairs of clusters in Fig. 3B, D, E and F, only sequence pairs where each sequence is from a separate cluster are used in the computation. As explained above, *mcorrPairGenome* and *mcorrViralGenome* both sort the CDS regions of the coronavirus genome into one continuous coding sequence, therefore input XMFA files must be prepared as described in the previous section *“Measurement of correlation coefficient of synonymous substitutions”*.

### Fitting procedure for correlation profiles and model selection

The fitting procedure used here is largely described in (30, 31). We used the LMFIT python package version 0.9.7 (85) to fit the analytical form of *P*(*l*) appearing in Eq. S2 (link to package here: https://lmfit.github.io/lmfit-py/). To infer recombination parameters, we fit the data with Eq. S2 by either varying the parameters *θ_s_*, *ϕ_s_*, and fixing 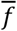 to be the length of the genome, or by varying all three parameters. The former fit is the “template-switching model”, which assumes only complete RNA templates are exchanged during template-switching events, and the latter is the “fragment-incorporation model”, which assumes that template-switching events can involve incomplete RNA templates (i.e., fragments). Both are described in the main text, and all data shown are the results of fitting with the template-switching model except for the parameters shown in Table S2. To distinguish between profiles with distinct signatures of recombination and instances of either no recombination or unclear signals of recombination, we compared the fits from the template-switching and fragment-incorporation models to what we refer to as the ‘null-recombination’ model. In the null-recombination model, we simply set *c*_*s*1_, = *c*_*s*2_ = 0 in Eq. S2, yielding 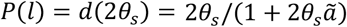. This gives a correlation profile independent of *l*, which is fit by averaging the measured values of *P*(*l*), yielding a single parameter: θ_*s*_. We then perform model selection using all three of these fits. To do this, we first use LMFIT to calculate the Akaike information criterion (AIC) value for each model, and then we compute the Akaike weight for each model, which is loosely considered to be the probability that a given model best predicts the data (86, 87). The Akaike weight for model *j* is calculated as:

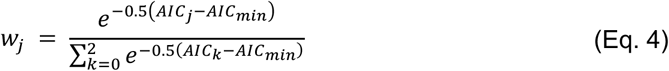

where the denominator sums over all three models and *AIC_min_* is the minimum of the AIC values. Finally, we calculate the evidence ratio for a given model *j* compared to a model *m*, which represents the likelihood of a given model being favored over another with respect to minimization of Kullback-Leibler discrepancy (87), by taking: *w_j_/w_m_.* In all tables (except Table S2), we interpret instances where the evidence ratio is *w_t_/w_n_* ≤ 100 for the template-switching model (t) compared to the null-recombination model (*n*) to mean that there is a lack of evidence for recombination. For Table S2, we fit correlation profiles with the fragment-incorporation model and interpret cases where *w_f_/w_n_* ≤ 100 to mean there is a lack of evidence for recombination (where denotes the Akaike weight for the fragment-incorporation model). For all tables, if there is a lack of evidence for recombination (i.e., model selection determines the null recombination model best fits the data), we simply calculate *θ_sample_* using *d_sample_* and the standard population genetic expression for heterozygosity (63) given by:

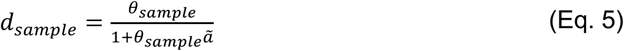

where *ã* = 4/3. Fitting and calculation of AIC values for correlation profiles averaged over all sequence pairs was performed with *mcorr-viral-fit* (developed for this study). Fitting and model selection with AIC for the correlation profiles of individual sequence pairs (as was done in Fig. 3) was performed using *mcorr-pair-fit*, which was also developed for this study.

### Clustering using pairwise synonymous diversity

For this study, we developed a CLI program *calcKsPair* to take XMFA files of whole genome alignments and measure pairwise synonymous diversity (*d_pair_*) across the genome using Eq. 2. Here, the fraction of synonymous substitutions for the third-position sites of codons is computed across all CDS regions of the genome, as opposed to computing *d_s_* for each gene and taking the average of these *d_s_* values. Similar to *mcorrLDGenome*, when we filtered out gene alignments with >10% gaps, *FilterGaps* was used with the flag “fill-gaps” on; this is a requirement for *calcKsPair* as well, which also sorts the CDS regions by the position they appear in the genome into one continuous sequence of codons. We then used the program *mcorr-dm* (developed in (31) and can be found here: https://github.com/kussell-lab/mcorr-clustering) to collect outputs into a square distance matrix for use with standard clustering algorithms. Lastly, we clustered sequences using the unweighted pair group method with arithmetic mean (UPGMA, also known as ‘average linkage’ (57)) using *d_pair_* as the distance metric. Clustering and dendrogram visualization was performed using the *scipy* and *matplotlib* python packages.

### Phylogenetic inference by maximum likelihood

We used *Augur* (88) with MAFFT (89) to align the 191 SARS-like coronavirus whole genome sequences to a SARS-CoV-1 reference genome (GenBank accession number MK062183.1). *IQ-Tree* was then used to build a maximum-likelihood tree using the generalized time reversible (GTR) model. The tree shown in Fig. S8 was visualized with Dendroscope3 (90) and re-rooted on the BtKY72 sequence (cluster 11 in Fig. 3A-B).

### Visualization of recombination network graphs

Visualization of recombination networks in Fig. 3D and S7 were performed with the python package *graph-tool* version 2.43 (91).

### Estimate of SARS-CoV-2 mutation rate

We estimated the SARS-CoV-2 mutation rate (to compute the mean coalescence time of sequence pairs shown in Fig. 4E) using the latest global subsampling of SARS-CoV-2 sequences from GenBank performed by *Nextstrain* (52–54). This dataset was downloaded on January 14^th^, 2022 from the *Nextstrain* site for SARS-CoV-2 (https://nextstrain.org/ncov/open/global). *Augur* (53) was used to build a tree with *IQ-TREE* (64) and *TreeTime* (54) was used to estimate the mutation rate.

## Acknowledgments and Funding Sources

This work was supported by NIH grant R01-GM-097356 (to E.K.) and grant 20/1041 from the Health Research Council of New Zealand (to O.K.S). Asher Preska Steinberg is a Simons Foundation Awardee of the Life Sciences Research Foundation. We gratefully acknowledge the New York University (NYU) high performance computing cluster for resources, and its staff for technical support.

## Supplementary Information

### Supplementary Information Text

#### Coalescent-based population genetics model with recombination

The coalescent-based model with recombination was originally described with application in bacteria in (1). In bacteria, the fragment-incorporation model is appropriate as it corresponds to uptake of DNA fragments and incorporation into the genome via homologous recombination. Here we give a brief summary of the model equations necessary for fitting correlation profiles and derive the functional form of the template switching model which is specific for RNA viruses. Model derivation and full description of parameters are given in the “Supplementary Notes” of (1).

The sample’s measured synonymous diversity (*d_sample_*) is a linear combination of the pool diversity [*d*(*Θ_p_*)] (from recombination), and the sample’s clonal diversity [*d*(*Θ*_s_)]:

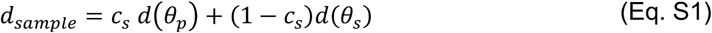

where *c_s_* is the sample’s recombination coverage and *d*(*θ*) is the standard population genetic expression for heterozygosity (2) (see table below). We abbreviate *θ_pool_*, *ϕ_pool_*, *θ_sample_, ϕ_sample_* as *θ_p_, ϕ_p_, θ_s_, ϕ_s_*, respectively. The predicted form of the correlation profile is:

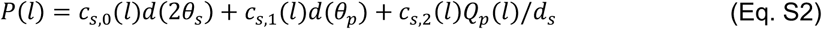

where the functional forms for *c_s,i_*(*l*) (*i* = 0,1,2) and *Q_p_*(*l*) are given in the table below. The expressions in the table correspond to the fragment-incorporation model, and the next paragraph describes how the expressions for the template-switching model are obtained.

The general expressions for *c_s_* and *c_s,i_*(*l*) are derived in (1), and can be specialized to different kinds of recombination (e.g., template switching or fragment incorporation) by computing the total rate (*r*) at which a single locus (for *c_s_*), or a pair of loci a distance *l* apart (for *c_si_*(*l*)), is affected by recombination per generation per sequence. This computation is illustrated for the two viral recombination models in Fig. S1, where *γ* is the recombination rate per site per sequence per generation. When computing the recombination rate for a pair of loci, recombination can affect either only one locus (‘one-site’ recombination which occurs at a rate of *r*_1_) or both loci (‘two-site’ recombination which occurs at a rate of *r*_2_). For one-site recombination, in the template-switching model the recombination breakpoint must land between the two sites, hence the rate of events is *r*_1_ = *γl*. In the fragment-incorporation model, the breakpoint must additionally land no further than the fragment size 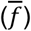 away from the locus in question; thus, *r*_1_ = *γl* for 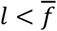, or 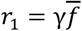 for 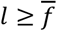. For a two-site recombination, in the template-switching model the breakpoint must occur either before the first locus or after the second locus, but not between the loci; hence the rate of events is *r*_2_ = *γ*(*L* – *l*I), where *L* is the genome length. In the fragment-incorporation model, for two-site events the fragment must cover both sites, hence the rate of events is 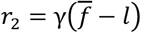 for 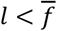, or *r*_2_ = 0 for 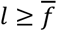. Summing the one- and two-site recombination rates yields the total rate of events that impact a single locus, which is *r* = *γL* for template-switching and 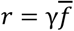 for fragment incorporation. We see therefore that the substitution 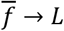 in the fragment-incorporation model (given in the table below) yields the template-switching model.

We can re-express Eq. S2 in terms of the three parameters *θ_s_, ϕ_s_*, and 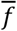, determined by fitting the profiles (as described in (1)). Subsequently, the pool parameters are obtained using the following relations 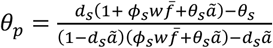 and *ϕ_p_* = *θ_p_ϕ_s_/θ_s_*; see table below for values of constants *ã* and *w*. For the template-switching model, we substitute *L* for 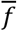, and fix the value of this parameter to be the genome length in question; the template-switching model thus involves only two free parameters.

| Term | Expression | Description |
| --- | --- | --- |
| $\mu$ | | Mutation rate<br>(per nt per generation) |
| $\gamma$ | | Recombination rate<br>(per nt per generation) |
| $\bar{f}$ | | Mean fragment size (nt) of<br>homologous recombination |
| $L$ | | Genome length (nt) |
| $l$ | | Distance (nt) between two loci |
| $\bar{T}_s$ or $\bar{T}_p$ | | Mean coalescence time for a pair<br>from the sample or pool. |
| $\theta_s$ or $\theta_p$ | $\theta_s = 2\mu T_s, \theta_p = 2\mu T_p$ | Mutational divergence of the<br>sample or pool |
| $\phi_s$ or $\phi_p$ | $\phi_s = 2\gamma T_s, \phi_p = 2\gamma T_p$ | Recombinational divergence of<br>the sample or pool |
| The following constants are used below: $\tilde{a} = 4/3$ and $w = 2/3$. | | |
| $d(\theta)$ | $\frac{\theta}{1 + \theta \tilde{a}}$ | Synonymous diversity<br>(heterozygosity) for divergence $\theta$ |
| $c_s$ | $\frac{\phi_s w \bar{f}}{1 + \theta_s \tilde{a} + \phi_s w \bar{f}}$ | Recombination coverage of the<br>sample |
| $c_{s,1}(l)$ | $\frac{2\phi_s w l}{1 + 2\theta_s \tilde{a} + \phi_s w (\bar{f} + l)}$ | Probability that the most recent<br>event was a recombination<br>affecting only one locus |
| $c_{s,2}(l)$ | $\frac{\phi_s w (\bar{f} - l)}{1 + 2\theta_s \tilde{a} + \phi_s w (\bar{f} + l)}$ | Probability that the most recent<br>event was a recombination<br>affecting both loci |
| $c_{s,0}(l)$ | $1 - c_1 - c_2$ | Probability that the most recent<br>event at either locus was not a<br>recombination |
| $Q_p(l)$ | $2 \left( \frac{\theta_p}{1 + \theta_p \tilde{a}} \right)^2 \left( \frac{1 + \theta_p \tilde{a} + \phi_p w l}{1 + 2\theta_p \tilde{a} + 2\phi_p w l} \right)$ | Probability of a difference at both<br>loci in the pool |

#### Clonal structure of samples and the residual clonality effect size

A key aspect of our analysis is the partitioning of the sample’s diversity (*d_sample_*) into two parts according to Eq. S1: The clonal diversity, *d_clonal_* = *d*(*θ_sample_*), generated by mutations that occurred within the sample since its coalescence, and the pool diversity, *d_pool_* = *d*(*θ_pool_*), generated by recombination events that bring different alleles into the sample from external sources. From the model fit, we obtain *θ_pool_* which determines *d_pool_* via Eq. S1. Then, by comparing *d_pool_* and *d_sample_* we determine whether any detectable clonal signal remains in the data. Specifically, if a sample has recombined extensively with the pool, the coverage *c_s_* will be very close to 100%, and *d_sample_* and *d_pool_* may be statistically indistinguishable depending on the uncertainty in our determination of *d_sample_*. In contrast, a statistically significant difference between *d_pool_* and *d_sample_* indicates that the fraction of the genome that has not been affected by recombination can be distinguished from the fraction of the genome which comes from the pool, enabling detection of a clonal signal within the sample. In this case, we can apply Eq. S1 to infer the value of *d_clonal_*, from which we obtain the sample’s mutational divergence *θ_sample_*.

We assess the statistical significance of the clonal signal by defining the residual clonality (RC) effect size as follows:

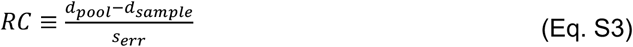

where 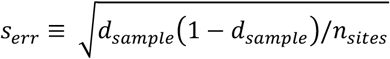, and *n_sites_* is the average number of positions over which *d_sample_* is calculated; thus, 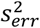 is the expected variance in the estimation of *d_sample_*. The RC effect size determines how large the difference between the pool and sample diversity is in units of the diversity measurement’s standard deviation; as such, *RC* takes the same general form as the widely used effect size statistic known as Cohen’s *d* statistic (3, 4). Values of RC < 1 indicate that insufficient clonal signal remains in the data, and therefore measurement of *θ_sample_* is expected to be unreliable. In Fig. S6-7, we plotted *c_sample_* and *θ_sample_* versus *RC* for cluster pairs in the SL-CoV dataset.

#### Estimation of sample’s recombination coverage from distributions of zero-SNP blocks

We consider a pair of sequences, and we measure the ‘substitution profile’ (*σ_i_*) at each four-fold degenerate, third-codon position *i* for the given pair, where *σ_i_* = 1 for a difference and *σ_i_* = 0 for identity. We assume that consecutive blocks of *σ_i_* = 0 (i.e., zero-SNP blocks) beyond a certain length are due to clonal portions that have not recombined (noting that sequencing error rates in the SL-CoV data can be estimated to be ~1e-5 (56)). Our prediction is that this clonal fraction is 1 - *c_sample_*. Within the genomic portions that have recombined, we still expect to find zero-SNP blocks, and the distribution of these block sizes is determined by *d_pool_.*

We define the ‘0-portion’ of the genome to be the set of all sites where *σ_i_* = 0, and the ‘1-portion’ as those sites where *σ_i_* = 1. The size of the 0-portion is ≈ 1 - *d_sample_* and, correspondingly, the size of the 1-portion is ≈ *d_sample_.* The ‘0-portion’ contains recombined and clonal sites. However, based on our observations that *θ_pool_* » *θ_sample_*, we expect that the 1-portion is almost entirely composed of recombined sites. We can therefore use the 0-portion to gain insights about the fraction of recombined sites (i.e., *c_sample_*) by predicting the distribution of zero-SNP block lengths which is expected in parts of the genome that have recombined with the pool, and determining how this prediction deviates from the empirical distribution. We expect that uncharacteristically long zero-SNP blocks are due to clonality, and we can visualize these by looking at the sliding window averages 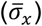 in Fig. S11A-B for the sequence pairs with lower values of *c_sample_*. For Fig. S11C, with high *c_sample_*, we see these uncharacteristically long zero-SNP blocks are no longer visually noticeable as their block length decreases to the limit of or below the sliding window size, and the clonal portion of the genome is fragmented by many recombined portions (we note that this pattern suggests a recombination mechanism which includes some degree of fragment incorporation in addition to template switching). Therefore, as we expect that zero-SNP blocks above a certain length are due to clonality, the block length *x\** above which we begin to observe block lengths longer than predicted is the clonal fraction of the 0-portion, which we denote this genomic fraction as *F*\*. Thus, consistency with the inference of *c_sample_* from the coalescent model is seen if we observe *c_sample_* ≈ 1 - *F*\*(1 – *d_sample_*).

In regions that have recombined with the pool, our null hypothesis is that the zero-SNP blocks will have a geometric distribution, which can be approximated in the continuous limit as

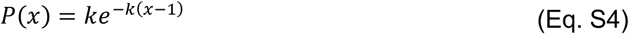

where *x* is the length of a zero-SNP block, and *k* is the probability of *σ_i_* = 1; given the recombined regions exhibit the pool’s diversity, we expect *k* ≈ *d_pool_*. The probability that a randomly chosen site for which *σ_i_* = 0 is part of a block of length *x* is given by:

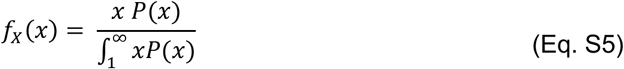

The associated cumulative distribution function (*F_X_*(*x*)) can be found by integrating Eq. S5:

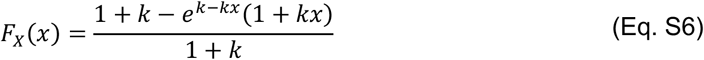

In Fig. S12, *F_X_*(*x*) is plotted along with the length-weighted empirical cumulative distribution functions 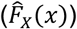 of zero-SNP block lengths for the three sequence pairs also shown in Fig. S11. As described in the Fig. S12 figure caption, we find that the values of *c_sample_* predicted from the coalescent model are similar to those predicted by looking at the value of *x* for which 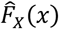 and *F_X_*(*x*) begin to deviate. We find that when fitting Eq. S6 to 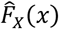 the two high coverage sequence pairs yield values of *k* close to *d_pool_* while *k* for the low coverage sequence pair (Rs4081 vs Rs4874) is lower than *d_pool_*, which may be a result of higher variance in the sampling of the pool diversity for pairs with low recombination coverage.

#### Simulations of heterogenous mutation rates using *PhastSim*

We used the “hypermutability” model of the *PhastSim* (5) package to simulate rate heterogeneity across genomic sites in SARS-like coronaviruses. *PhastSim* achieves this by allowing the user to specify that a certain proportion of sites have a specific mutation or transition rate (i.e., the transition from one specific nucleotide to another) be enhanced by some factor *μ*, as described in eq. 2 of ref. (5). One can then choose the proportion of positions with enhanced or “hypermutable” rates. The algorithm takes as input a tree and reference genome, then simulates sequence evolution across the phylogeny using a Gillespie algorithm. As input, we first generated a random tree with a minimum of 1000 tips using a Birth-Death model with the software package *ngesh* (6), resulting in a tree with 2018 tips. The NCBI SARS-CoV-2 reference genome (Table S5) was used as a reference genome. For non-hypermutable sites, the UNREST substitution model (7) was used with transition rates inferred for SARS-CoV-2 (8). Recent work has suggested that for SARS-CoV-2, *G* → *U* and *C* → *U* transitions have elevated rates (8), with some portion of these sites being hypermutable. Furthermore, this work shows that *G* and *C* sites account for ~6.5% and ~11.4% of synonymous sites, respectively, and that their transition rates are on average ~5-100-fold greater than other transition rates, with some of these sites being hypermutable. Therefore, in the simulations in Fig. S6, we first conducted simulations with the average estimated heterogeneous transition rates inferred for SARS-CoV-2 (8) without hypermutable sites (labeled “heterogeneous”). As it is unclear what portion of *G* and *C* sites could be hypermutable, we simulated extreme scenarios where the proportion of sites with elevated rates were 15-20% (which would account for all *G* and *C* sites), and their enhanced mutation rates from *μ* = 10 - 100. To calculate the sliding window averages and correlation profiles shown in Fig. S6, we created a FASTA file as outputs from the simulations, then generated an XMFA file as described in Materials and Methods. The sliding window averages in Fig. S6A-B were calculated using measurements of the substitution profile at each site (measured by *mcorrLDGenome* as described in Materials and Methods subsection “Measurement of correlation coefficient of synonymous substitutions”). The correlation profiles across the genome in Fig. S6C-D were calculated and fit as described in Materials and Methods.

**Fig. S1.**
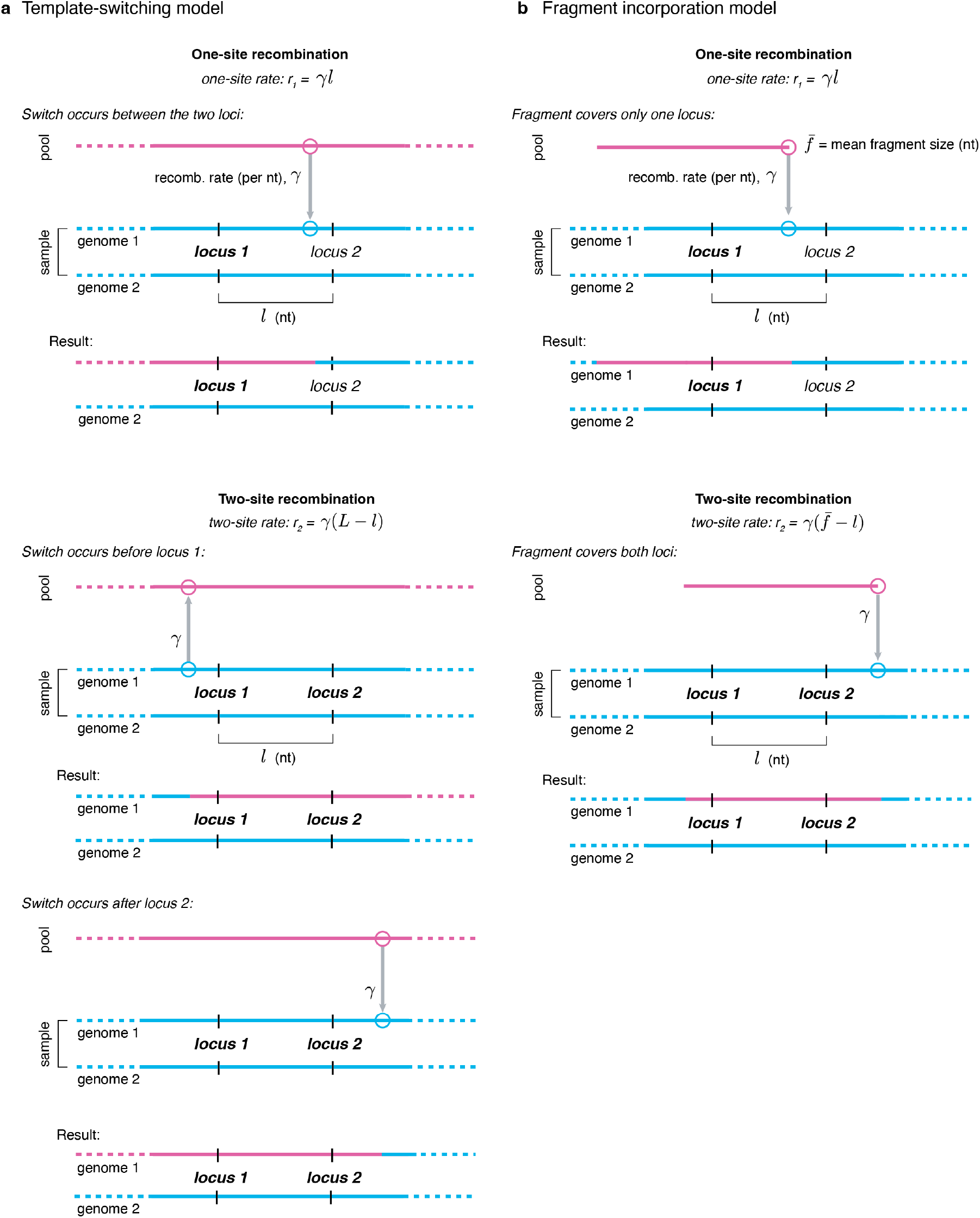
Schematic depicting different types of recombination events in the “template-switching model” and the “fragment-incorporation” model. One- and two-site recombination events are depicted for the template-switching model (A) and the fragment-incorporation model (B).

**Fig. S2.**
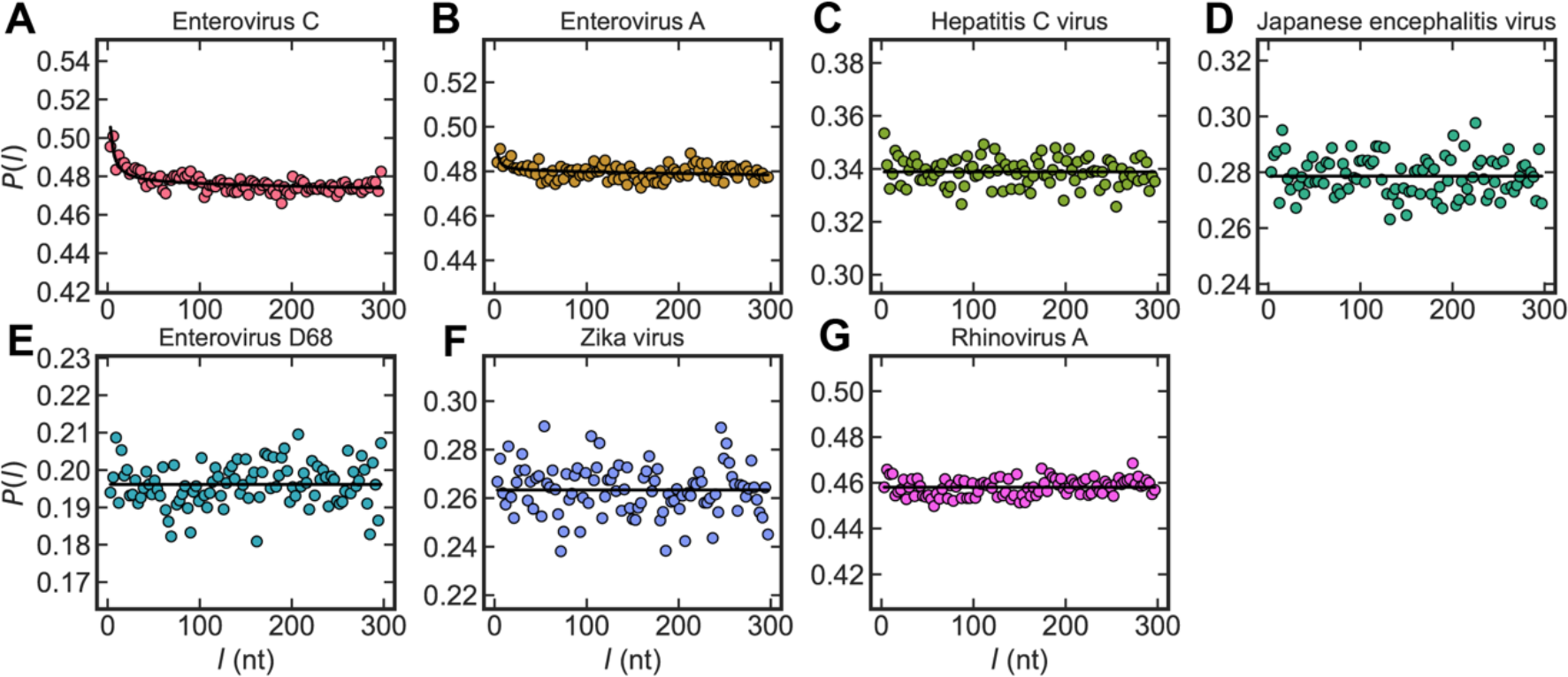
Correlation profiles of synonymous substitutions for seven additional (+)ssRNA viruses. Markers correspond to the correlation profile *P*(*l*) for a given separation distance *l* (given in nucleotides, *nt).* The fit is shown as a solid line, and was performed under the assumption that only complete RNA strands can be exchanged during template-switching (i.e., we used the “template-switching model” described in the Main Text and Materials and Methods). Model selection was performed with the Akaike Information Criterion (AIC) to determine if a coalescent model with or without recombination best fit the data (see Materials and Methods for details). Parameters of homologous recombination are given in Table S1.

**Fig. S3.**
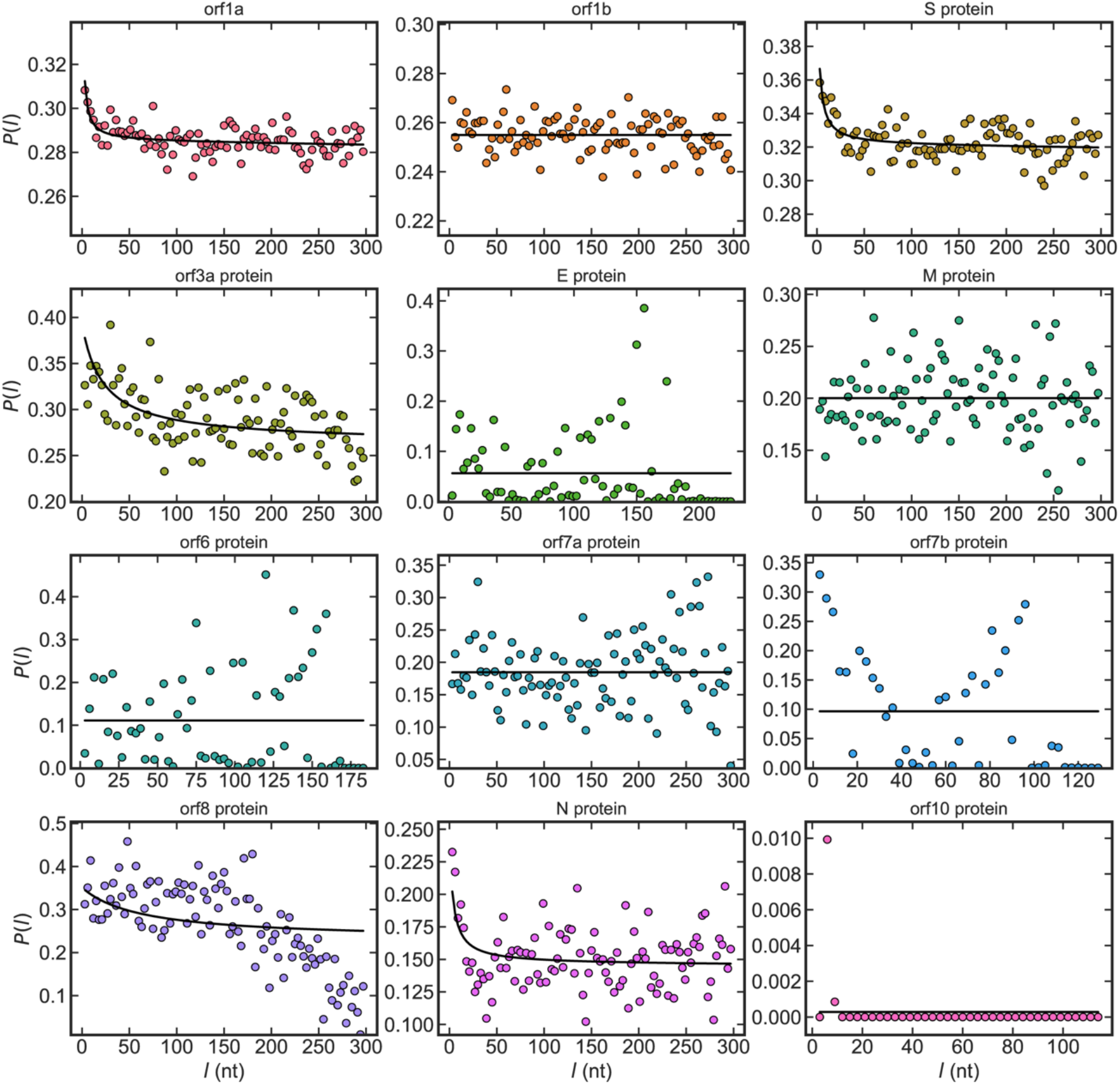
Correlation profiles of synonymous substitutions for all genes in the SARS-like betacoronavirus genome. Markers correspond to the correlation profile *P*(*l*) for a given separation distance *l* (given in nucleotides, *nt*). The fit is shown as a solid line, and was performed under the assumption that only complete RNA strands can be exchanged during template-switching (i.e., we used the “template-switching model” described in the Main Text and Materials and Methods). Model selection was performed with the Akaike Information Criterion (AIC) to determine if a coalescent model with or without recombination best fit the data (see Materials and Methods for details). Parameters of homologous recombination are given in Table S3.

**Fig. S4.**
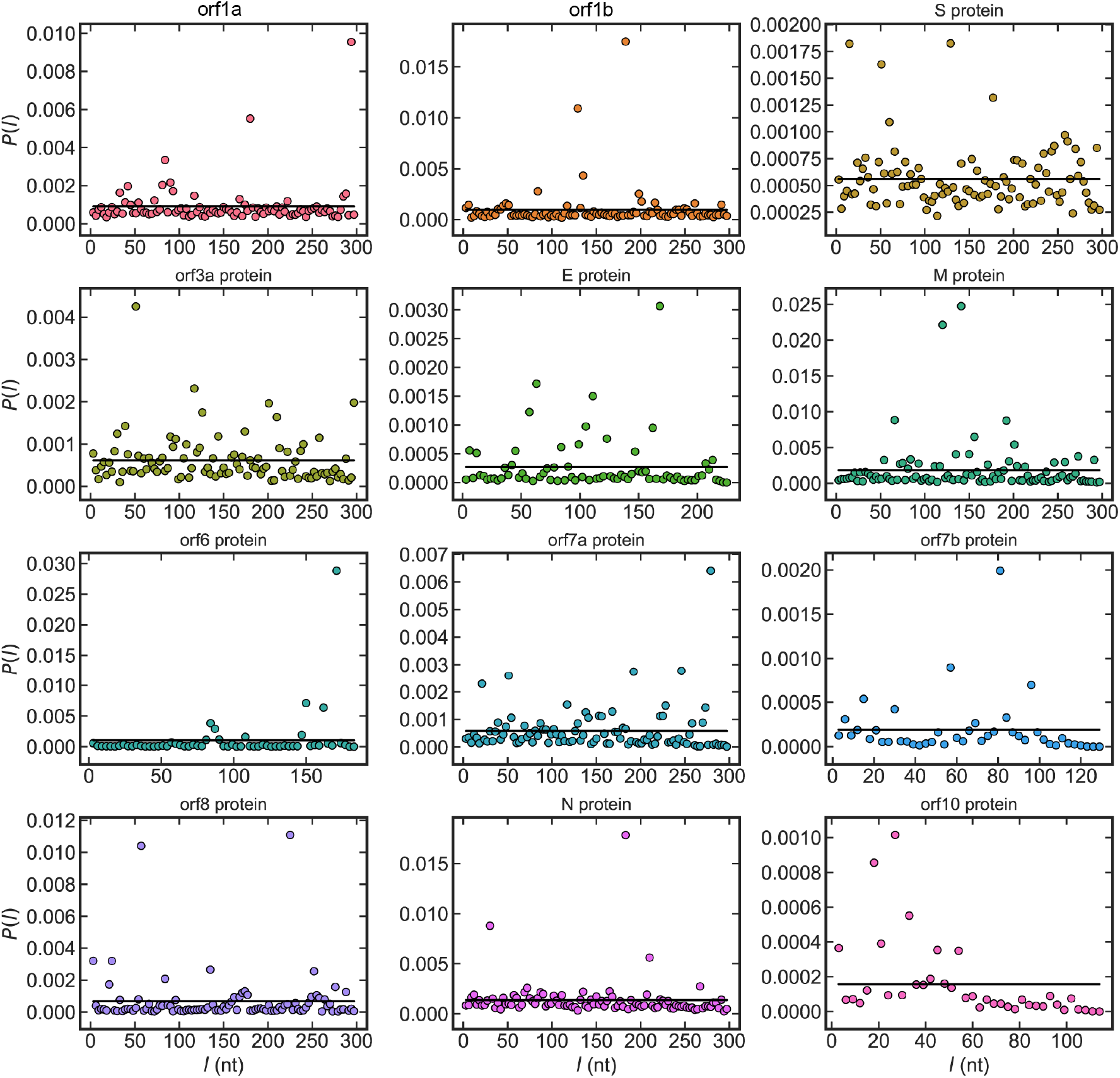
Correlation profiles of synonymous substitutions for all genes in the SARS-CoV-2 genome calculated for 444,145 SARS-CoV-2 sequences. Markers correspond to the correlation profile *P*(*l*) for a given separation distance *l* (given in nucleotides, *nt*). The fit is shown as a solid line, and was performed under the assumption that only complete RNA strands can be exchanged during template-switching (i.e., we used the “template-switching model” described in the Main Text and Materials and Methods). Model selection was performed with the Akaike Information Criterion (AIC) to determine if a coalescent model with or without recombination best fit the data (see Materials and Methods for details). Parameters of homologous recombination are given in Table S4.

**Fig. S5.**
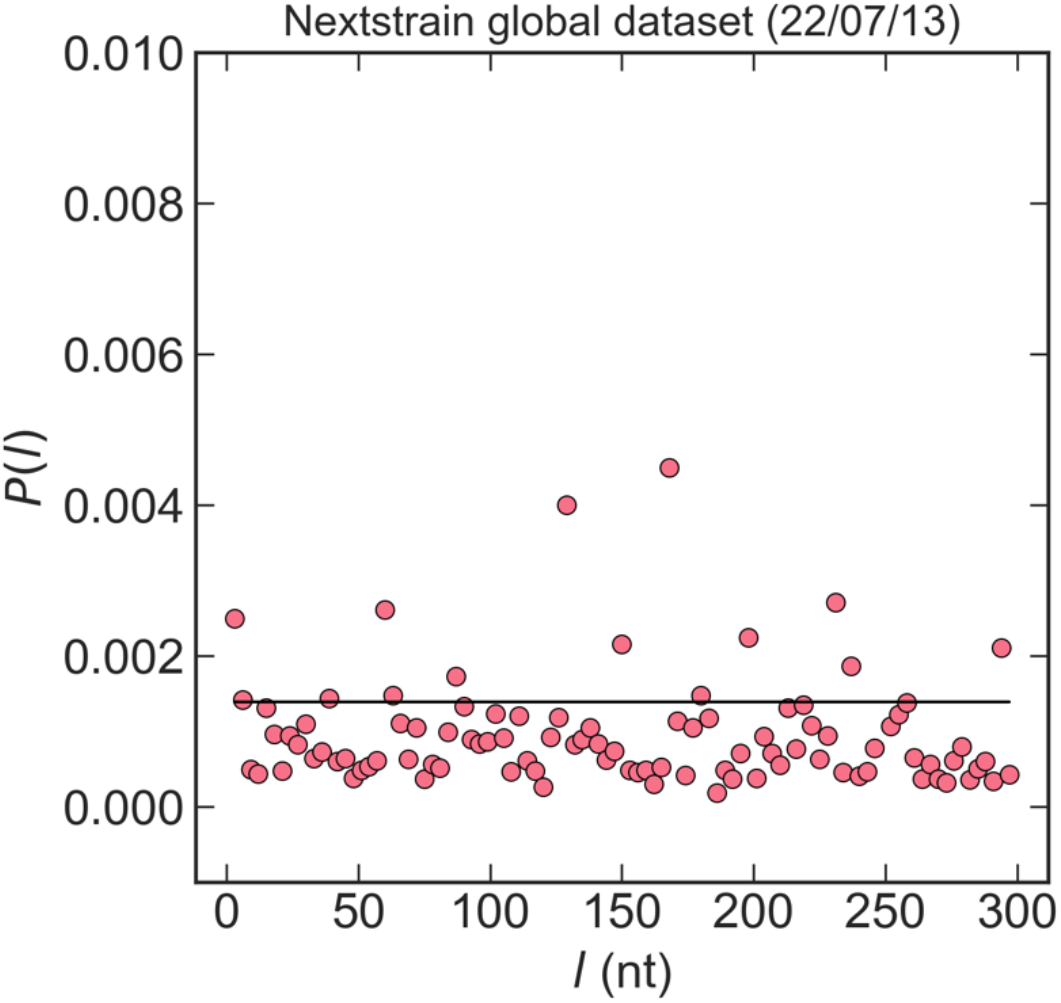
Correlation profile of synonymous substitutions across the whole genome of SARS-CoV-2 calculated using the latest *Nextstrain* global subsampling over the last six months (date of analysis was July 13^th^, 2022 and included 2,860 sequences). Markers correspond to the correlation profile *P*(*l*) for a given separation distance *l* (given in nucleotides, *nt*). Model selection was performed with the Akaike Information Criterion (AIC) to determine if a coalescent model with or without recombination best fit the data (see Materials and Methods for details). As the AIC values indicated that the coalescent model without recombination was the best fit for the data (the Akaike weight of the “template-switching model” over that of the “zero-recombination model” was: *w_t_/w_z_* = 0.37), the fit shown above (as a solid line) is the fit to the coalescent model without recombination.

**Fig. S6.**
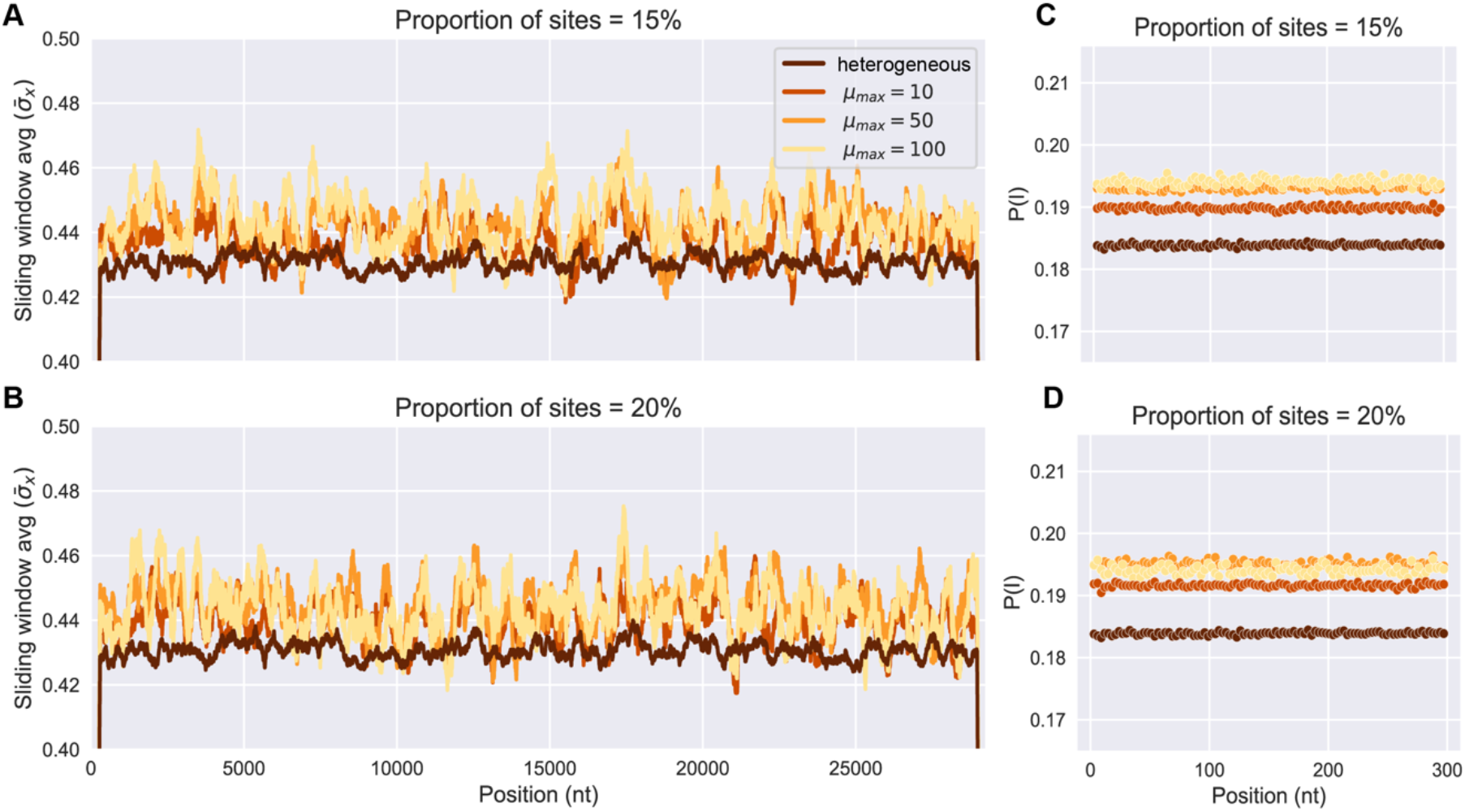
Results of simulations of heterogeneous mutation rates using the hypermutable model in the *PhastSim* package (55). (A-B) Sliding window averages of pairwise diversity 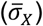 for third-position synonymous codons averaged across all simulated sequences. (C-D) Correlation profiles of synonymous substitutions calculated across the whole genome for all simulated sequences. We used model selection with the Akaike Information Criterion (AIC) to determine if a coalescent model with or without recombination was a better fit for the data. In all cases, AIC indicated that the data was best fit by the coalescent model without recombination (the highest ratio of the Akaike weight of the “template-switching model” over that of the “zero-recombination model” was: *w_t_/w_z_* = 3.7 × 10^-52^). The proportion of nucleotide positions which are “hypermutable” with enhanced transition rates is indicated in the titles of each figure subpanel. The enhanced transition rate (*μ_max_*) for these sites is indicated in the legend of subpanel *A.* For all other sites, transition rates are set to be the average estimated mutation rates for SARS-CoV-2 from SI ref. (8). All panels also include the results of simulations with the average estimated mutation rates and no hypermutable sites (labeled “heterogenous”). Additional details on simulations can be found in Supplementary Information.

**Fig. S7.**
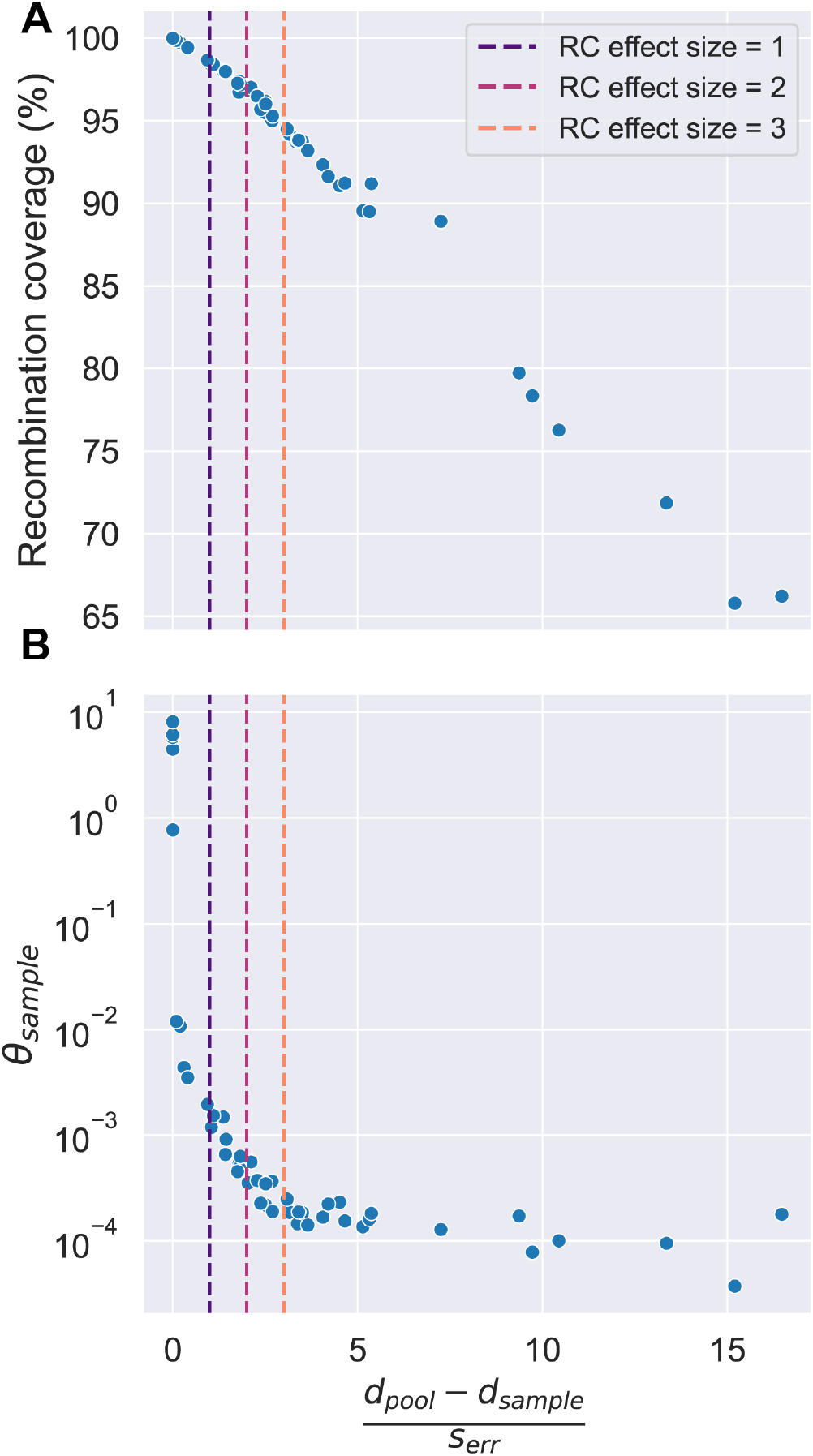
Dependence of the sample’s recombination coverage (A) and its mutational divergence (*θ_sample_*) (B) on the residual clonality (RC) effect size for pairs of clusters from the SARS-like coronavirus clusters from main text Fig. 3A-B. The RC effect size is calculated as described by Eq. S3 of Methods. Recombination coverage is given as the percentage of recombined sites in the sample, *θ_sample_* has units of *nt*^-1^, and the RC effect size is unitless.

**Fig. S8.**
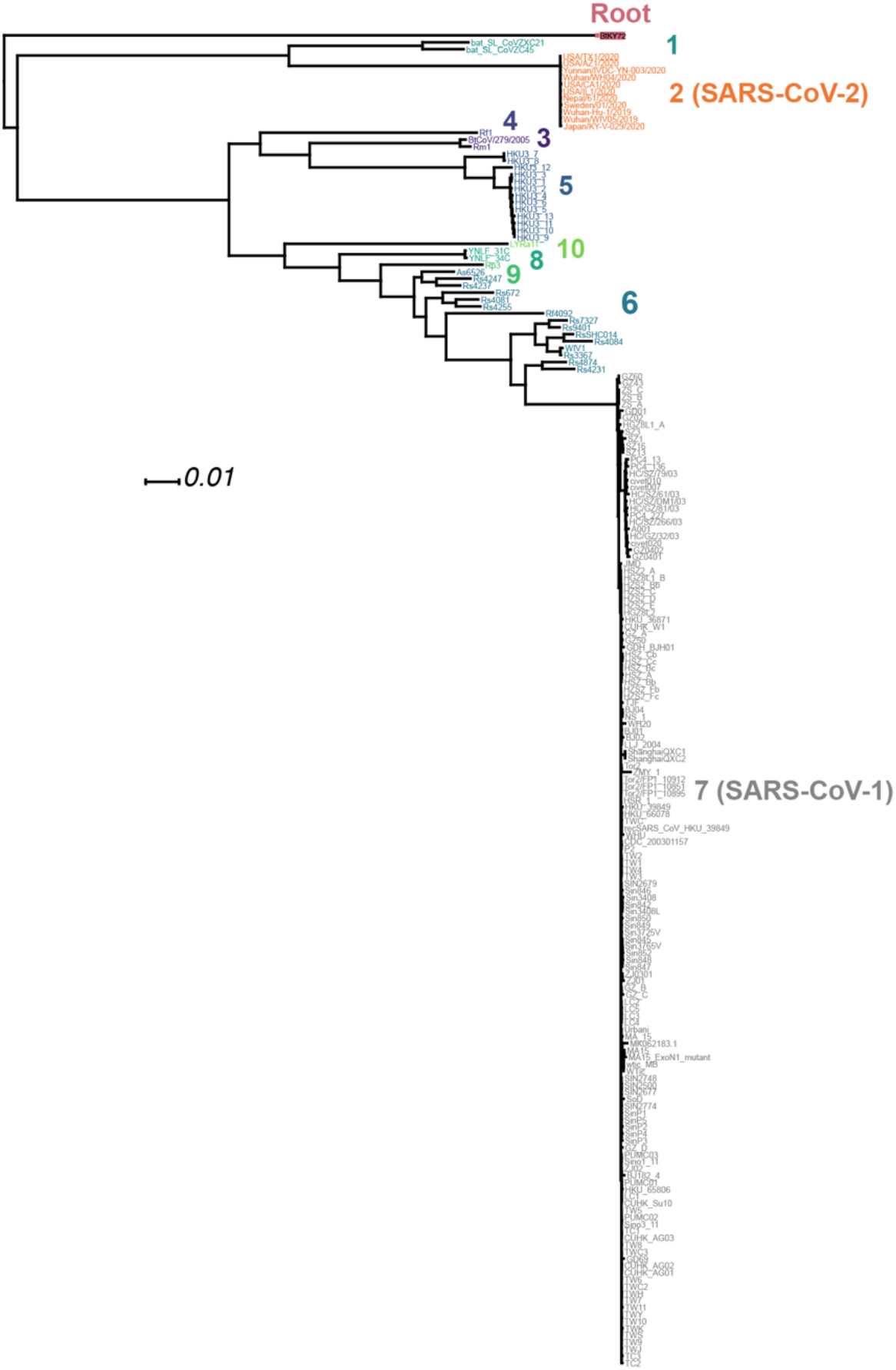
Phylogeny for the 191 strains shown in the dendrogram in main text Fig. 3A created by estimating the maximum-likelihood phylogeny with *IQ-Tree.* Color scheme is the same as Fig. 3A-B. Branch lengths and scale bar measure nucleotide divergence. Tips are strain names. Details on phylogenetic inference and visualization in Materials and Methods.

**Fig. S9.**
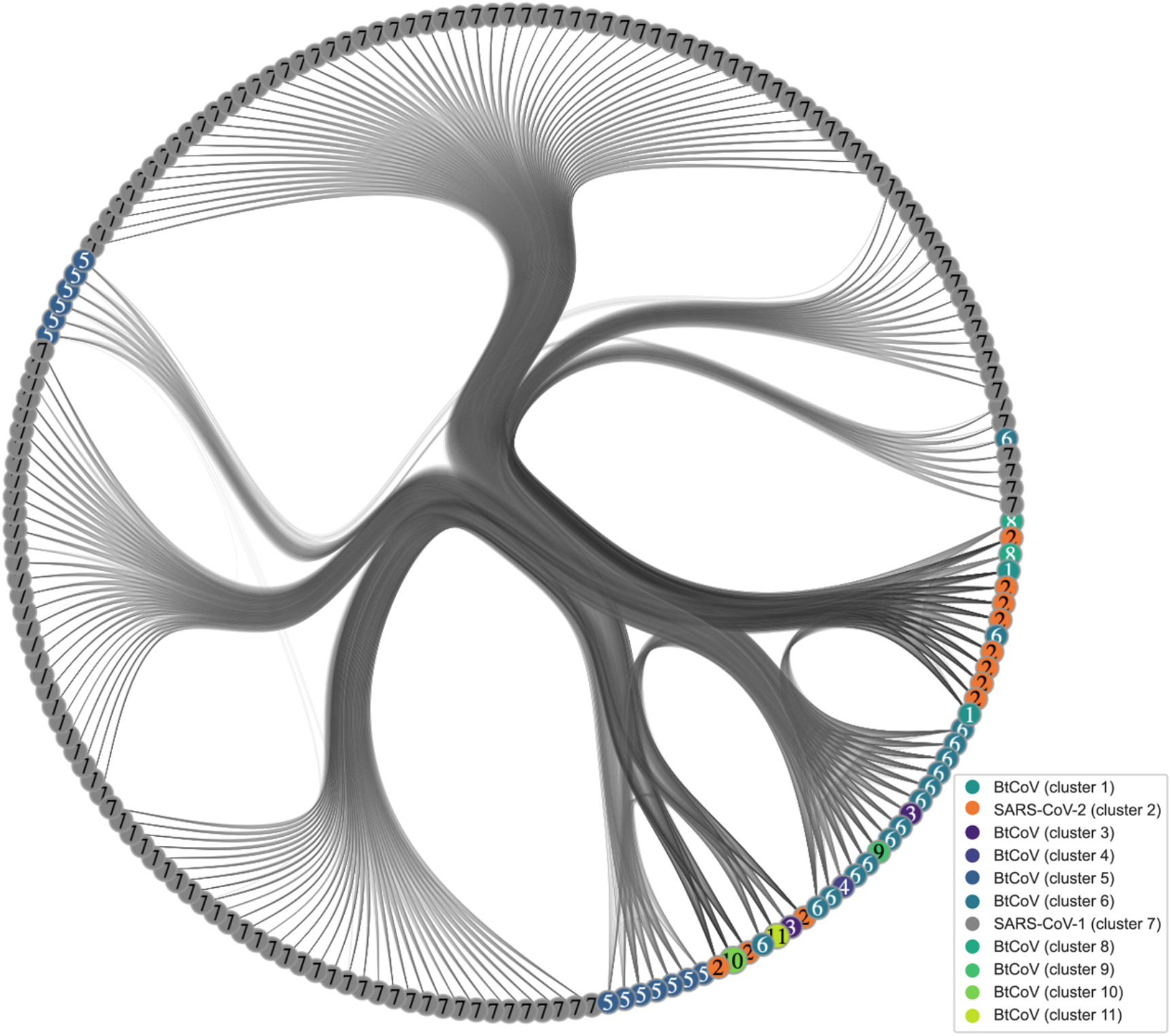
Recombination network graph for the 191 strains shown in the dendrogram in main text Fig. 3A. The heatmap shown in Fig. 3B corresponds to this network. The nodes are strains, and the color scheme of the nodes corresponds to the 11 flat clusters created by cutting the dendrogram at *d_s_* = 0.09 in Fig. 3A. The edges connecting nodes indicate that the pair of sequences has recombined with a shared pool, determined by computing correlation profiles across the whole genome and fitting these profiles to the model. Edge shading is to allow for better visualization of edges. Numbers on nodes correspond to sequence clusters. In the legend, clusters composed of SARS-like coronaviruses from bats are abbreviated as “BtCoV”. The same fitting procedure was used as Figs. 1-3 of the main text (see Materials and Methods). If model selection suggested the profile was better fit with the null-recombination model, no edge was assigned for the sequence pair.

**Fig. S10.**
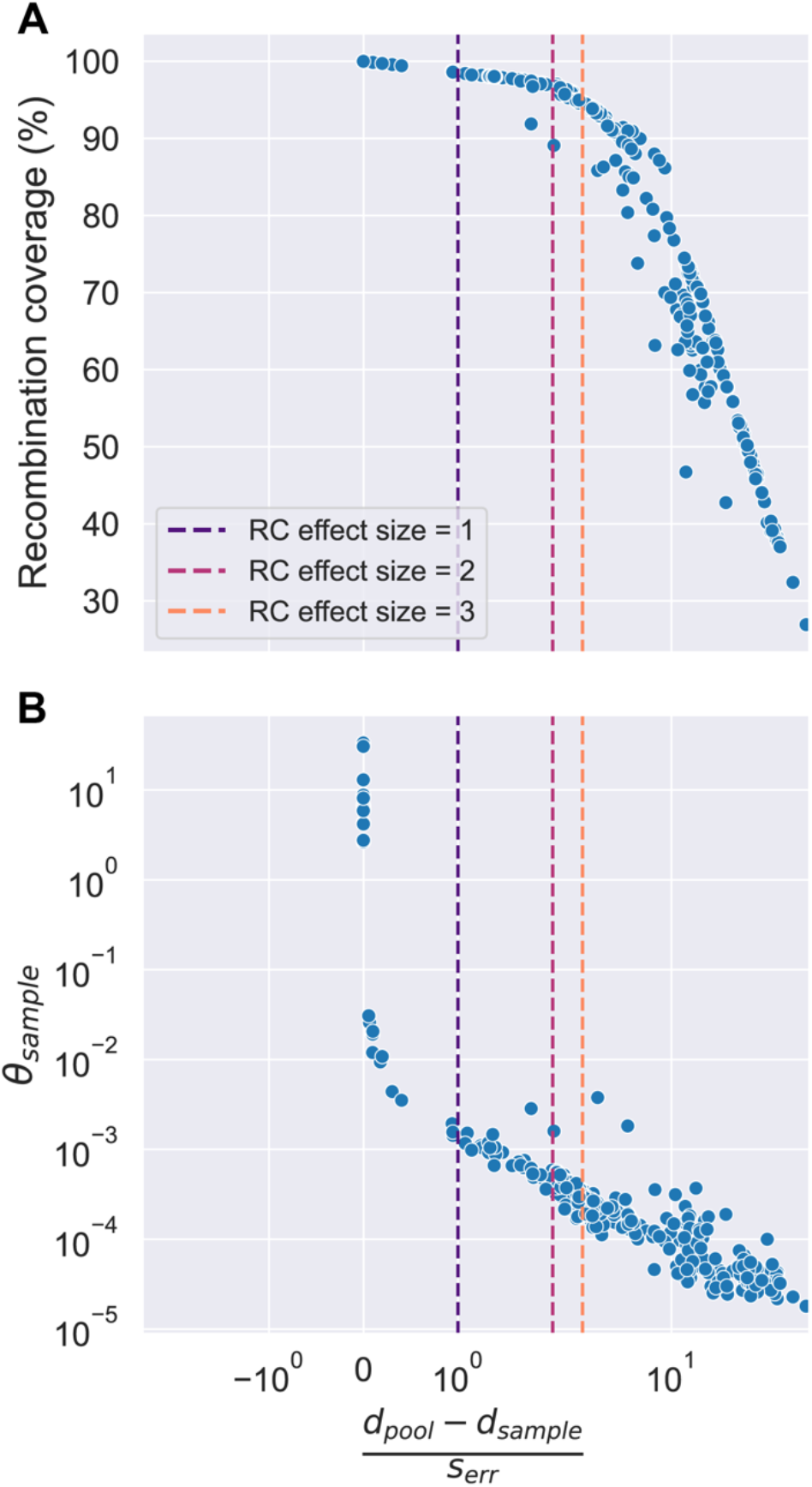
Dependence of the sample’s recombination coverage (A) and its mutational divergence (*θ_sample_*) (B) on residual clonality (RC) effect size evaluated for pairs of clusters from the 27 SARS-like coronavirus clusters from main text Fig. 3E-F. The RC effect size is calculated as described by Eq. S3 of Methods. Recombination coverage is given as the percentage of recombined sites in the sample, *θ_sample_* has units of *nt*^-1^, and the RC effect size is unitless.

**Fig. S11.**
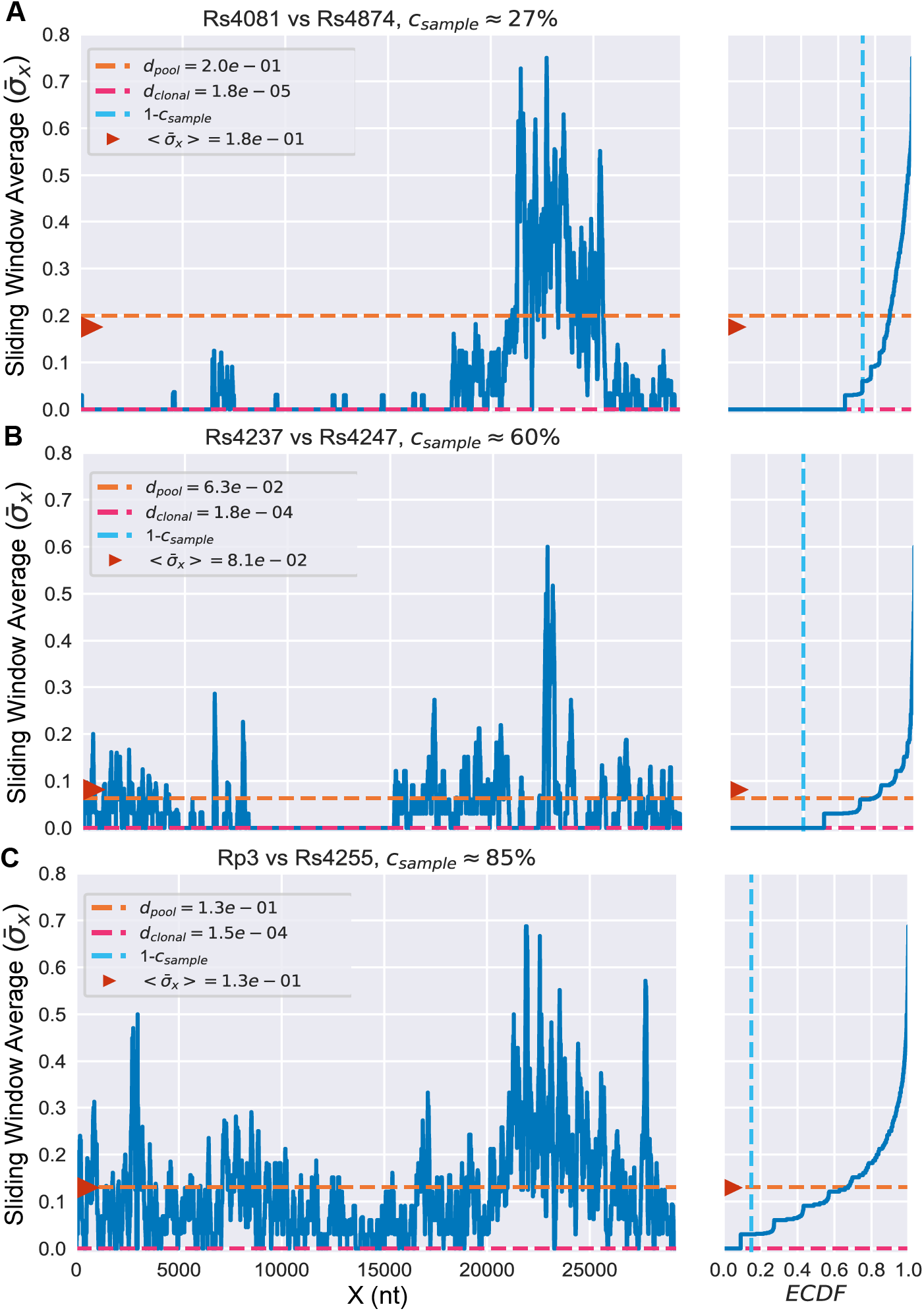
Sliding window averages of pairwise diversity 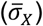 for third-position synonymous codons calculated for three pairs of SARS-like coronaviruses (SL-CoVs). Plots on the left show 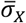 plotted against genomic position in nucleotides (nt). Plots on the right show empirical cumulative distribution functions (ECDFs) of 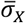. Dotted lines indicate values of parameters inferred from the template-switching model. The red arrows indicate the average value of all 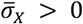, which is close to the inferred value of *d_pool_*, as would be expected if regions with 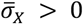 are recombined in from the pool.

**Fig. S12.**
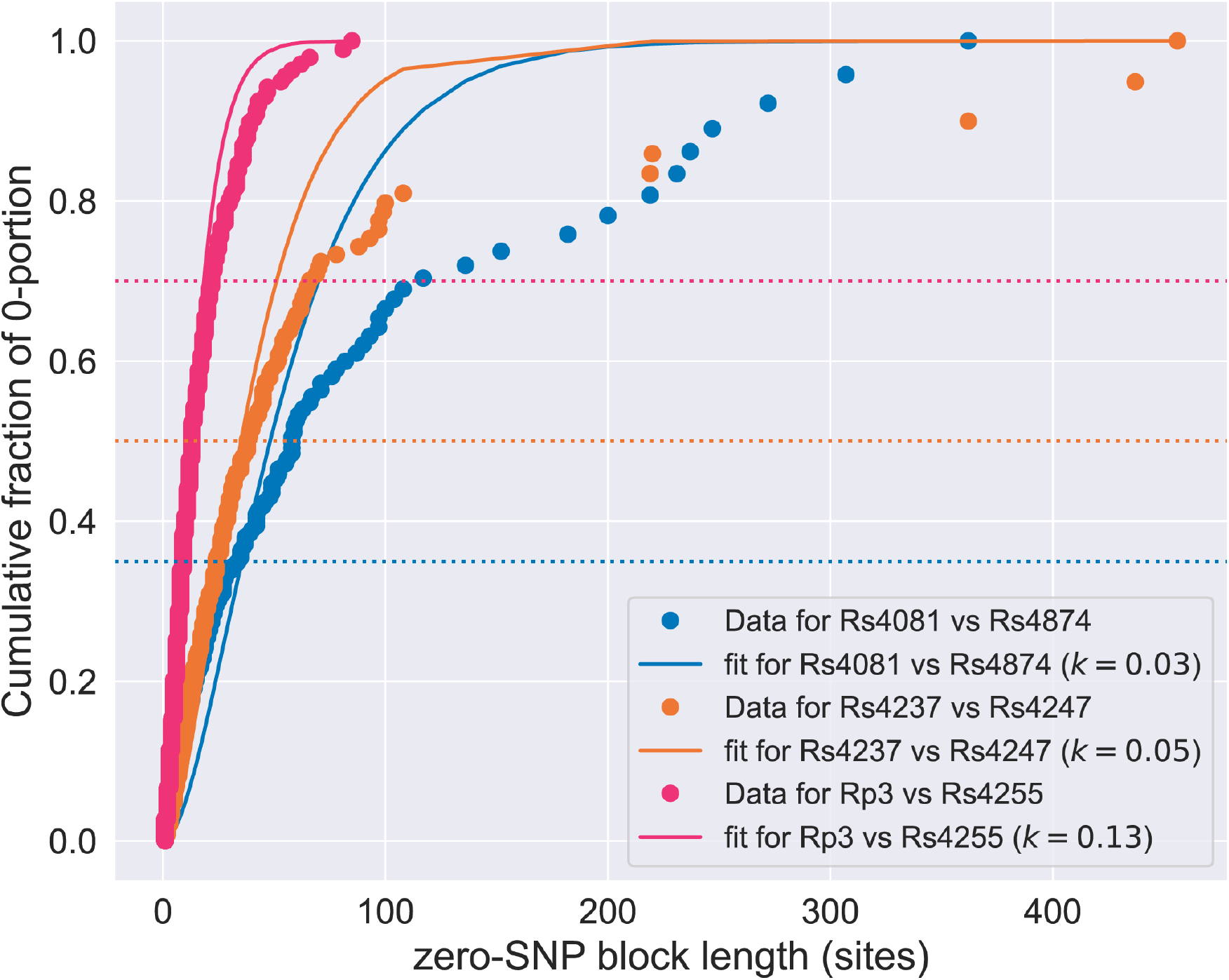
Length-weighted distributions of zero-SNP block lengths for the sequence pairs shown in Fig. S11. Length-weighted empirical cumulative distribution functions 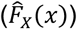 of zero-SNP block lengths (consecutive number of sites for which the synonymous diversity given by Eq. 2 is zero) are plotted as markers. To find the corresponding cumulative distribution function (*F_X_*(*x*); plotted as solid lines), Eq. S6 was fit to 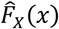 by using the Trust Region Reflective algorithm and allowing *k* to vary between 0 and ∞. As described in the section of the Supplemental Information titled *Estimation of sample’s recombination coverage from distributions of zero-SNP blocks*, the fraction at which 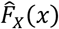 and *F_X_*(*x*) begin to diverge can be used to estimate *c_sample_* by use of the relationship: *c_sample_* ~ 1 - *F*\*(1 – *d_sample_*). Dotted lines depict the approximate fraction at which this divergence occurs. If we estimate that *F*\* ~ 0.65, 0.5, and 0.3 for Rs4081 vs Rs4874, Rs4237 vs Rs4247, and Rp3 vs Rs4255, respectively, then we can estimate that *c_sample_* ~ 40%, 50%, and 70% for these pairs. As shown in Fig. S11, the template-switching model predicts that *c_sample_* ~ 27%, 60%, and 85% for these sequence pairs.

**Table S1.**
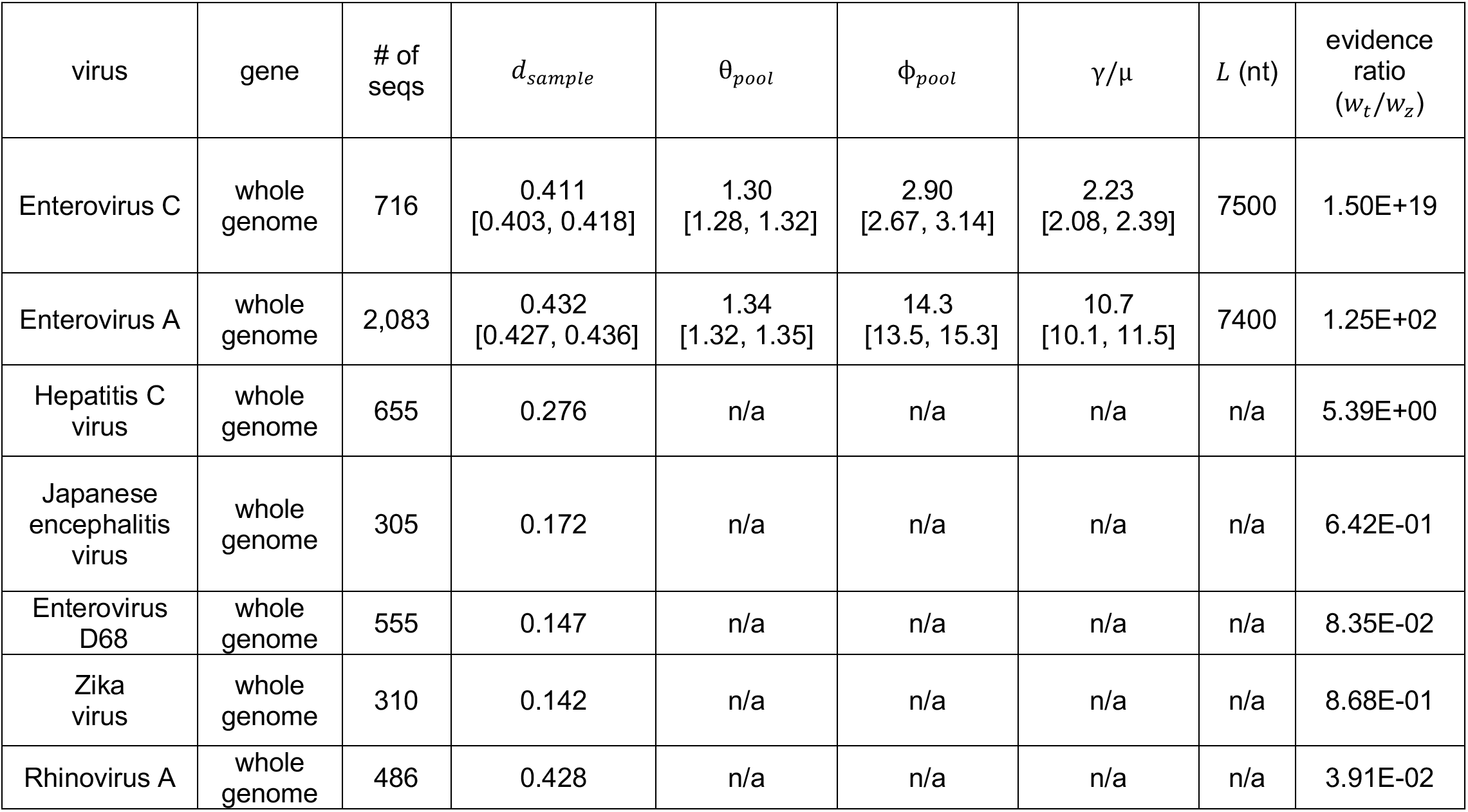
Parameters of homologous recombination for viruses shown in Fig. S2. If the entire genome was fit, the gene is listed as “full genome”. Parameters are given as the values inferred from the data, followed by the 95% bootstrap CI in square brackets (see Materials and Methods for calculation). We used model selection with AIC to determine if the profile was better fit with a coalescent model with or without recombination (see Materials and Methods). For those profiles which were better fit with the coalescent model with recombination, we assumed a “fixed fragment size” (i.e., only complete RNA strands are exchanged during template-switching). For those profiles better fit by the model without recombination, coverage was set to c = 0 and no bootstrapping was performed. *L* ≡ length of the RNA strand exchanged during template-switching, *w_f_/w_z_* is the Akaike weight of the recombination model with fixed fragment size (*w_f_*) over the weight of the model without recombination (*w_z_*; see Materials and Methods). Other parameters are described in the main text.

| virus | gene | # of seqs | $d_{sample}$ | $\theta_{pool}$ | $\phi_{pool}$ | $\gamma/\mu$ | $L$ (nt) | evidence ratio ($w_t/w_z$) |
| --- | --- | --- | --- | --- | --- | --- | --- | --- |
| Enterovirus C | whole genome | 716 | 0.411<br>[0.403, 0.418] | 1.30<br>[1.28, 1.32] | 2.90<br>[2.67, 3.14] | 2.23<br>[2.08, 2.39] | 7500 | 1.50E+19 |
| Enterovirus A | whole genome | 2,083 | 0.432<br>[0.427, 0.436] | 1.34<br>[1.32, 1.35] | 14.3<br>[13.5, 15.3] | 10.7<br>[10.1, 11.5] | 7400 | 1.25E+02 |
| Hepatitis C virus | whole genome | 655 | 0.276 | n/a | n/a | n/a | n/a | 5.39E+00 |
| Japanese encephalitis virus | whole genome | 305 | 0.172 | n/a | n/a | n/a | n/a | 6.42E-01 |
| Enterovirus D68 | whole genome | 555 | 0.147 | n/a | n/a | n/a | n/a | 8.35E-02 |
| Zika virus | whole genome | 310 | 0.142 | n/a | n/a | n/a | n/a | 8.68E-01 |
| Rhinovirus A | whole genome | 486 | 0.428 | n/a | n/a | n/a | n/a | 3.91E-02 |

**Table S2.**
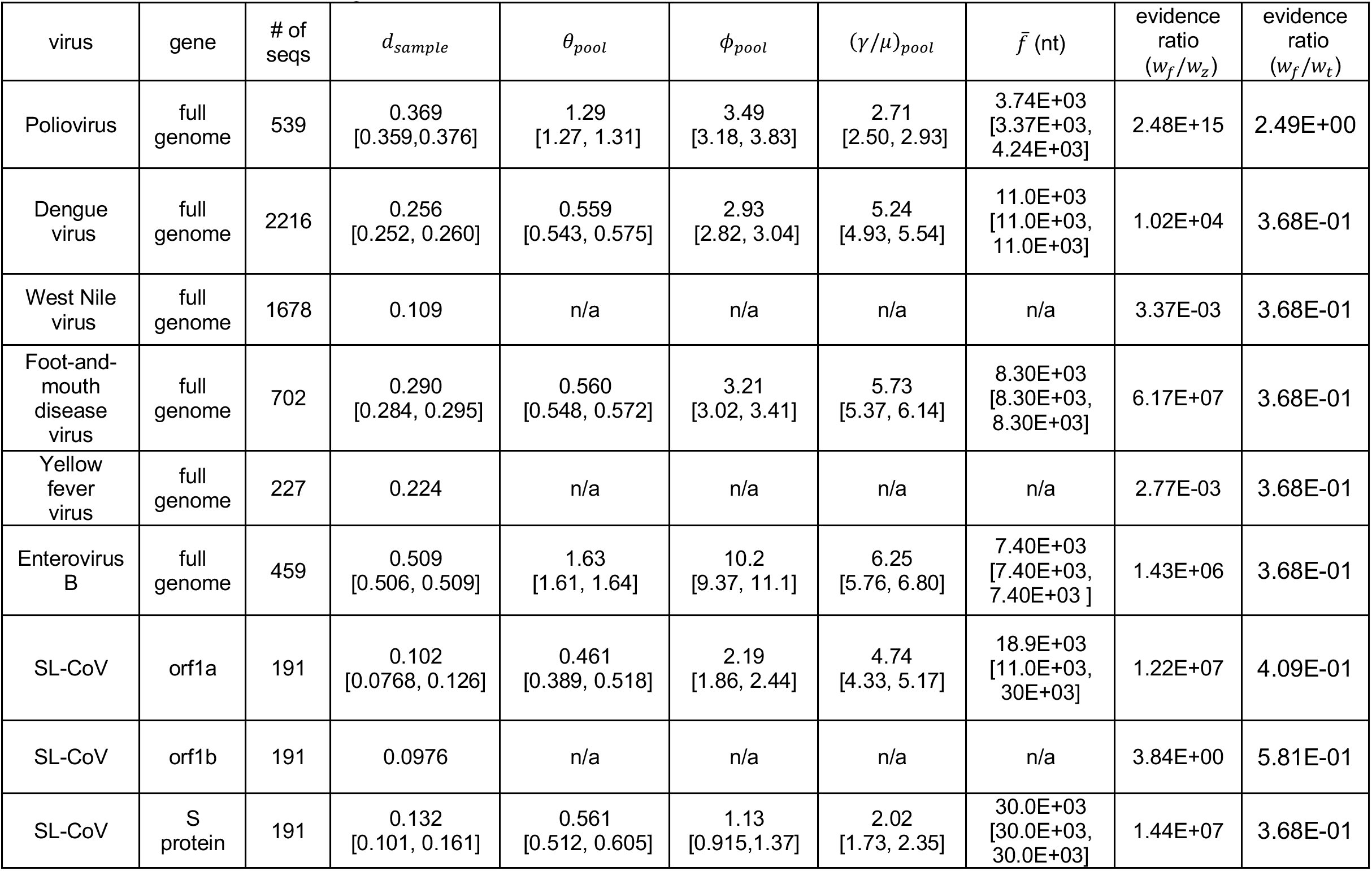
Parameters of homologous recombination from fitting viruses discussed in main text with the recombination model under the assumption that template-switching events can occur with full RNA templates and fragments (i.e., the “fragment-incorporation model” described in the Main Text and Materials and Methods). If the entire genome was fit, the gene is listed as “full genome”. Parameters are given as the values inferred from the data, followed by the 95% bootstrap CI in square brackets (see Materials and Methods for calculation). We used model selection with AIC to determine if the profile was better fit with a coalescent model with or without recombination (see Materials and Methods). For those profiles better fit by the model without recombination, coverage was set to c = 0 and no bootstrapping was performed. 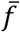 ≡ the mean size of a recombined fragment, *w_f_/w_z_* is the Akaike weight of the fragment-incorporation model (*w_f_*) over the weight of the model without recombination (*w_z_*), and (*w_f_ /w_t_*) is the Akaike weight of the recombination model with fragment-incorporation model over that of the template-switching model (*w_t_*). Other parameters are described in the main text. See Materials and Methods for more details on models and fitting.

**Table S3.** Parameters of homologous recombination for each gene of 192 SARS-like coronavirus whole genome sequences (corresponding correlation profiles shown in Fig. S3). Parameters are given as the values inferred from the data, followed by the 95% bootstrap CI in square brackets (see Materials and Methods for calculation). We used model selection with AIC to determine if the profile was better fit with a coalescent model with or without recombination (see Materials and Methods). For those profiles better fit by the model without recombination, coverage was set to c = 0 and no bootstrapping was performed. For the CDS region of orf8 protein, while the evidence ratio (*w_t_/w_z_*) favored the coalescent model with recombination and this fit is displayed in Figure S2, this did not appear to adequately describe the correlation profile. Thus, we simply give *d_sample_* and *θ_sample_* estimated from the model without recombination. *L* ≡ length of RNA strand exchanged during template-switching (fixed at the genome length), *w_t_/w_z_* is the Akaike weight of the template-switching model (*w_t_*) over the weight of the model without recombination (*w_z_*; see Materials and Methods). Other parameters are described in the main text.

| virus | gene | $d_{sample}$ | $\theta_{pool}$ | $\phi_{pool}$ | $(\gamma/\mu)_{pool}$ | $L$ (nt) | evidence ratio ($w_t/w_z$) |
| --- | --- | --- | --- | --- | --- | --- | --- |
| SL-CoV | orf1a | 0.102<br>[0.0768, 0.126] | 0.460<br>[0.400, 0.516] | 2.04<br>[1.74, 2.34] | 4.45<br>[4.03, 4.84] | 3.00E+04 | 2.98E+07 |
| SL-CoV | orf1b | 0.0976 | n/a | n/a | n/a | n/a | 6.61E+00 |
| SL-CoV | S protein | 0.132<br>[0.101, 0.161] | 0.561<br>[0.512, 0.605] | 1.13<br>[0.915, 1.37] | 2.02<br>[1.73, 2.35] | 3.00E+04 | 3.91E+07 |
| SL-CoV | orf3a protein | 0.0892<br>[0.0671, 0.112] | 0.413<br>[0.274, 0.477] | 7.13E-02<br>[3.75E-03, 1.97E-01] | 0.172<br>[0.0132, 0.440] | 3.00E+04 | 1.02E+05 |
| SL-CoV | E protein | 0.0278 | n/a | n/a | n/a | n/a | 7.69E-01 |
| SL-CoV | M protein | 0.0677 | n/a | n/a | n/a | n/a | 3.62E-01 |
| SL-CoV | orf6 protein | 0.0416 | n/a | n/a | n/a | n/a | 3.68E-01 |
| SL-CoV | orf7a protein | 0.0708 | n/a | n/a | n/a | n/a | 3.55E-01 |
| SL-CoV | orf7b protein | 0.0492 | n/a | n/a | n/a | n/a | 6.19E+00 |
| SL-CoV | orf8 protein | 0.230 | n/a | n/a | n/a | n/a | 1.76E+04 |
| SL-CoV | N protein | 0.0461<br>[0.0329, 0.0593] | 0.182<br>[0.111, 0.265] | 2.86E-01<br>[2.40E-04, 4.11E-01] | 1.57<br>[0.00153, 2.07] | 3.00E+04 | 2.01E+03 |
| SL-CoV | orf10 protein | 0.00399 | n/a | n/a | n/a | n/a | 1.63E-16 |

**Table S4.** Parameters of homologous recombination for each gene of SARS-CoV-2 inferred using 444,145 SARS-CoV-2 sequences (corresponding correlation profiles shown in Fig. S4). We used model selection with AIC to determine if the profile was better fit with a coalescent model with or without recombination (see Materials and Methods). In this case all correlation profiles were better fit with the model without recombination. *L* ≡ length of RNA strand exchanged during template-switching (fixed at the genome length), c is the recombination coverage given as a percentage of genomic sites, *w_t_/w_z_* is the Akaike weight of the template-switching model (*w_t_*) over the weight of the model without recombination (*w_z_*; see Materials and Methods). Other parameters are described in the main text.

| virus | gene | $d_{sample}$ | $\theta_{sample}$ | $\phi_{sample}$ | $\theta_{pool}$ | $\phi_{pool}$ | $(\gamma/\mu)_{pool}$ | $L$ (nt) | evidence ratio ($w_t/w_z$) |
| --- | --- | --- | --- | --- | --- | --- | --- | --- | --- |
| SARS-CoV-2 | orf1a | 9.96E-04 | 9.97E-04 | n/a | n/a | n/a | n/a | n/a | 2.71E-01 |
| SARS-CoV-2 | orf1b | 1.24E-03 | 1.24E-03 | n/a | n/a | n/a | n/a | n/a | 1.48E-01 |
| SARS-CoV-2 | S protein | 5.97E-04 | 5.97E-04 | n/a | n/a | n/a | n/a | n/a | 1.67E-01 |
| SARS-CoV-2 | orf3a protein | 7.59E-04 | 7.59E-04 | n/a | n/a | n/a | n/a | n/a | 1.70E-02 |
| SARS-CoV-2 | E protein | 4.02E-04 | 4.02E-04 | n/a | n/a | n/a | n/a | n/a | 1.95E-02 |
| SARS-CoV-2 | M protein | 1.83E-03 | 1.84E-03 | n/a | n/a | n/a | n/a | n/a | 3.68E-01 |
| SARS-CoV-2 | orf6 protein | 5.92E-04 | 5.93E-04 | n/a | n/a | n/a | n/a | n/a | 3.67E-01 |
| SARS-CoV-2 | orf7a protein | 6.75E-04 | 6.76E-04 | n/a | n/a | n/a | n/a | n/a | 1.94E-01 |
| SARS-CoV-2 | orf7b protein | 4.52E-04 | 4.52E-04 | n/a | n/a | n/a | n/a | n/a | 1.22E-05 |
| SARS-CoV-2 | orf8 protein | 1.86E-03 | 1.86E-03 | n/a | n/a | n/a | n/a | n/a | 1.02E-10 |
| SARS-CoV-2 | N protein | 1.03E-03 | 1.03E-03 | n/a | n/a | n/a | n/a | n/a | 3.76E-01 |
| SARS-CoV-2 | orf10 protein | 3.81E-04 | 3.81E-04 | n/a | n/a | n/a | n/a | n/a | 6.15E-07 |

**Table S5.** Genbank accession numbers for reference genomes used in this study.

| virus | Genbank accession |
| --- | --- |
| Poliovirus | AF462418.1 |
| Dengue virus | NC_001474.2 |
| West nile virus | NC_001563.2 |
| Foot-and-mouth disease virus | NC_039210.1 |
| Yellow fever virus | NC_002031.1 |
| Enterovirus B | NC_038307.1 |
| SARS-CoV-2 | NC_045512.2 |
| Enterovirus C | NC_002058.3 |
| Enterovirus A | NC_038306.1 |
| Hepatitis C virus | NC_038882.1 |
| Japanese encephalitis virus | MT560941.1 |
| Enterovirus D68 | NC_038308.1 |
| Zika virus | NC_035889.1 |
| Rhinovirus A | NC_038311.1 |

